# Phase-Specific Antibiotic Resistance Mechanisms in an *Escherichia coli* B Strain

**DOI:** 10.1101/2025.06.30.662419

**Authors:** Manuel Terrazas-López, Vanessa Aitken, Tonya N. Zeczycki, Eda Koculi

**Affiliations:** Department of Chemistry and Biochemistry, The University of Texas at El Paso, El Paso, TX 79968, USA; Department of Biochemistry and Molecular Biology, The Brody School of Medicine at East Carolina University, Greenville, NC 27858, USA

## Abstract

The majority of antibiotics developed to date target the fast-growing phase of bacteria, which typically occurs during active infection and resembles the exponential growth phase of laboratory-grown cultures. However, many pathogenic bacteria in the human body occupy environments where nutrients are limited and persist in a low-metabolic state, mirroring the stationary phase observed in laboratory-grown cultures. Bacteria in this latter phase are generally more resistant to antibiotics and are often responsible for recurrent infections. A comprehensive understanding of metabolic processes in both the exponential and stationary phases of bacterial growth could facilitate the development of phase-specific antibiotics. In this study, we adopted an untargeted proteomics approach to establish similarities and differences in the exponential and stationary phase proteome landscapes in BLR (DE3) cells, an *Escherichia coli* B strain. Our findings are consistent with previous studies, which show that metabolic processes such as translation and ribosome biogenesis decrease during the stationary phase compared to the fast-growing exponential phase, whereas tricarboxylic acid cycle activity increases. Importantly, our data reveal that distinct antibiotic resistance pathways are activated during the exponential and stationary phases, underscoring the potential for developing phase-specific antibiotic therapies.

## Introduction

Pathogenic *Escherichia coli* (*E. coli*) infections cause about 950,000 deaths worldwide each year ^1^. In addition, various nonpathogenic *E. coli* strains are important components of the human gut microbiome, while others are used in laboratories as model systems to investigate bacterial metabolic pathways, discover and produce novel antimicrobials, and serve as factories for protein production ^2–5^. Identifying the distinct molecular pathways activated under various stress conditions in different strains of nonpathogenic *E. coli* could improve our understanding of the microbiome, enhance our ability to use *E. coli* for protein and antimicrobial production, and guide the design of novel and much-needed antibiotics against pathogenic *E. coli* and bacterial infections more generally. To elucidate the proteomic differences underlying bacterial growth phases, we compared the relative protein abundances between the exponential and stationary phases of the nonpathogenic *E. coli* B strain BLR (DE3) using liquid chromatography–tandem mass spectrometry (LC-MS/MS) and a label-free, untargeted proteomics approach.

Our data on the *E. coli* B strain BLR (DE3) reveal that distinct antibiotic resistance pathways are activated during the exponential and stationary phases. This finding is significant, as traditional antibiotics are primarily designed to target fast-growing bacterial cells, which resemble the exponential phase of laboratory-grown cultures and dominate during active infections ^6^. However, during chronic infections, bacterial cells often enter the stationary phase, during which their metabolic activity slows and differs markedly from that in active infections ^6, 7^. There is a notable lack of antibiotics effective against bacteria in the stationary phase of growth ^6^. Ideally, novel antibiotics should either target both phases of bacterial growth or be tailored to specific stages of infection. This approach would help avoid administering antibiotics when they are least effective and thereby reduce the risk of developing antibiotic-resistant bacteria. The identification of distinct antibiotic resistance pathways activated during the exponential and stationary phases of *E. coli* BLR (DE3) could inform the design of phase-specific antibiotics for pathogenic *E. coli* and other bacterial species more broadly.

The proteomic changes between the exponential and stationary phases of the *E. coli* B strain, REL606, have been previously investigated. In that study, two-dimensional (2D) electrophoresis coupled with quadrupole time-of-flight (Q-ToF) MS was used to provide insights into the proteome of the *E. coli* REL606 strain in the exponential and stationary phases. However, due to the lower protein separation capacity of 2D electrophoresis compared to an offline reversed-phase fractionation system, only about 16% of the proteome could be inferred ^8^. In contrast, we implemented a high-resolution, label-free LC-MS/MS workflow using offline reversed-phase fractionation and Orbitrap-based detection to achieve substantially deeper proteome coverage. This approach enabled the confident identification and quantification of low-abundance regulatory proteins and metabolic enzymes not captured in earlier datasets, allowing us to identify approximately 43% of the proteome ^8^. This significantly enhanced depth of analysis revealed distinct profiles in antibiotic resistance, metabolic regulation, and translational control that predominate in each phase and were not detected in the previous study ^8^.

## Materials and Methods

### Materials

*E. coli* BLR (DE3) cells, which are a recA^-^ derivative of BL21 cells, were purchased from Novagen. The iST-Preparation and iST-Fractionation kits used for peptide generation and offline fractionation were purchased from PreOmics. All chemicals and reagents were purchased in the highest grade possible and used without further purification. Nondiet P-40 (NP-40) was originally purchased from Sigma Aldrich. (Note: IGEPAL CA-630 (Sigma--Aldrich) can be used as a direct replacement for NP-40). All remaining chemicals and reagents were obtained from Fisher Bioreagents.

## Methods

### Sample preparation

Five colonies *of E. coli* BLR (DE3) (*n = 5* biological replicates) were separately cultured in 50 mL Luria-Bertani (LB) broth overnight at 37 °C with continuous shaking at 225 revolutions per minute (rpm). Subsequently, the cultures were scaled up to 200 mL and adjusted to an initial optical density at 600 nm (OD_600_) of 0.03. The cultures were then incubated under the same growth conditions in the same medium for 3 hours, reaching an OD_600_ of approximately 0.3, and for 22 hours, reaching an OD_600_ of approximately 5. The cells were pelleted by centrifugation at 4 °C at 5000 x *g* for 15 minutes. The resulting cell pellets were flash-frozen in liquid nitrogen prior to peptide isolation.

### LC-MS/MS for Label-Free Proteomics Peptide isolation

Low protein binding microcentrifuge tubes (Eppendorf Protein LoBind Tubes) were used for peptide isolation and purification steps. Biological replicates (n=5) for each growth condition were initially resuspended in bacterial lysis buffer (500 µL, 50 mM Tris, pH 7.5, 150 mM NaCl, and 0.1% NP-40). It should be noted that IGEPAL CA-630 (Sigma Aldrich) can be directly substituted for NP-40. HALT Protease Inhibitor Cocktail (1X final concentration) was added to the lysis buffer immediately before use. The cells were disrupted *via* sonication on ice (10 sec burst, 30% amplitude, total of 1 min sonication). After sonication, proteins in the cell lysate were subsequently precipitated using ice-cold methanol (3:1 v/v) at −20° C overnight and pelleted by centrifugation (5000 x g, 10 min). The precipitated protein pellets were washed with ice cold methanol (500 µL, x 2) and allowed to air dry before further use.

PreOmics iST and iST-Fractionation Add-on Kits were used according to the provided instructions to prepare mass spectrometry grade peptides. Briefly, the precipitated proteins were resuspended in the provided denaturation/alkylation buffer (150 µL) and heated at 95 °C for 10 min. After cooling to room temperature, freshly prepared digestion solution (50 µL) was added to each sample. Samples were incubated at 37 °C for 2 hours to ensure complete digestion before adding the provided stop solution (100 µL). Samples were cooled to room temperature prior to transferring to the provided solid phase extraction columns. After 2 min of centrifugation (room temperature, 500 x g), the cartridges were washed with wash 1 and 2 solutions (200 µL each, 2 min centrifugation, 500 x g) to remove hydrophobic and hydrophilic contaminations. For reverse-phase, pH-dependent fractionation, peptides were eluted from the cartridges using elution buffers 1-3 (150 µL, 2 min centrifugation, room temperature, 500 x g). Each elution was collected in separate low protein binding tube and dried under a stream of N_2_ to dryness. Purified, alkylated peptides were reconstituted in 30 µL of loading buffer (98:2, v/v water/acetonitrile with 0.1% formic acid). Peptide concentrations in each sample were determined fluorometrically (Pierce Quantitative Peptide Assay) and adjusted to 0.25 mg/mL with loading buffer.

### nLC-MS/MS for label-free proteomics

Peptides were analyzed were subjected to nano–liquid chromatography–mass spectrometry (nanoLC-MS/MS) analysis using an UltiMate 3000 RSLCnano system (Thermo Fisher Scientific) coupled to a Q Exactive Plus Hybrid Quadrupole-Orbitrap mass spectrometer (Thermo Fisher Scientific) via a nanoelectrospray ionization source. For each fraction, 1 μg (4 µL) of sample was first trapped on an Acclaim PepMap 100 (20 mm x 0.075 mm) trapping column (Thermo Fisher Scientific) at 98:2 v/v water/acetonitrile with 0.1% formic acid (5 µL/min, 6 min trapping). Analytical separation was performed over 120 min with an effective linear gradient of 4-35% acetonitrile plus 0.1% formic acid (300 nL/min) using an EASY-Spray PepMap RSLC C18 column (75 μm x2 50 mm Thermo Fisher Scientific) with a column temperature of 50 °C. MS1 was performed at 70,000 resolution, with an automatic gain control (AGC) target of 1 × 10^5^ ions and a maximum injection time (IT) of 100 ms. MS2 spectra were collected by data-dependent acquisition on the top 20 most abundant precursor ions with a charge > 1 per MS1 scan. Precursor ion isolation window was 1.5 mass/charge ratio (m/z), dynamic exclusion was 20 s, and the normalized collision energy was 32. MS2 scans were performed at 17,500 resolution, maximum IT of 60 ms, and AGC target of 5 × 10^4^ ions.

### Database searching and analysis

FragPipe (v 19.1) was used to for raw data analysis with default search parameters for LFQ-MBR workflows ^9^. Fractions for each biological sample were searched together as a single experiment. An initial open search against the canonical Uniprot *E. coli* BLR(DE3) reference proteome (UP000475070, accessed 6/2024) was used to identify potential post-translational modifications for inclusion in the LFQ-MBR workflow ^10^. Precursor m/z tolerance was set to −150 to 500 Da and fragment tolerance was ±20 ppm with 3 missed cleavages for Trp and Lys-C allowed. Peptide spectrum matches (PSMs) were validated using PeptideProphet and results were filtered at the ion, peptide, and protein level with a 1% false discovery rate (FDR).

Based on these initial searches, the following variable modifications were included in the LFQ-MBR analysis: oxidation (+15.5995 Da on Met), deamidation (+0.98401 Da on Gln and Asn), and fixed modification carbamodiomethyl (+57.025 Da on Cys). For LFQ-MBR analysis, data were search again searched against the canonical Uniprot *E. coli* BLR(DE3) reference proteome (UP000475070, accessed 6/2024). Precursor ion m/z tolerance was ±20 ppm with 3 missed cleavages for Trypsin/LysC allowed. The search results were filtered by a 1% FDR at the ion, peptide, and protein-level. PSMs were validated using Percolator and label free quantification was carried out using IonQuant Match between runs FDR rate at the ion level was set to 10% for the top 300 runs. Proteins with >95% probability of ID, >2 unique peptides, and in more than 80% of a sample group (i.e. 4/5 injections) were considered high confidence IDs and retained for analysis. Intensities were log_2_ transformed, quantile normalized within the entire sample run and relative abundances for low sampling proteins were determined via knn methods (15 neighbors) in Perseus ^11^.

### Thresholds and biological process identification

GO enrichment analysis (statistical overrepresentation test) using Panther GO was used to identify over- and under-represented GO terms. A log_2_ fold change threshold of ± 0.585 (fold changes of 1.5 and 1/1.5) in protein abundances during the exponential and stationary growth phases was used to identify biologically significant differences in protein abundances and was included in the GO enrichment analysis ^12, 13^. Fisher’s exact test (*p* < 0.05) was used to establish statistically significant over-or under-representation. Protein–protein interactions and additional biological processes, molecular functions, and cellular locations of proteins were identified using STRING analysis with *E. coli* K-12 as the reference organism ^14^. A subset of protein functions was further investigated using UniProt and EcoCyc ^15, 16^.

## Results and Discussion

### 493 proteins showed altered abundance between the exponential and stationary phases of *E. coli* BLR (DE3), while 1,326 proteins remained unchanged

From approximately 4,200 predicted protein-coding genes in *E. coli*, we identified 1,819 proteins with high confidence (Figure 1) ^8^. By comparison, only 651 proteins were identified in the *E. coli* B strain REL606 study ^8^. Our findings therefore provide novel insights into alterations in metabolic pathways and cellular processes between the exponential and stationary growth phases—insights that were not attainable in the previous study due to the limited number of detected proteins ^8^.

**Figure 1.**
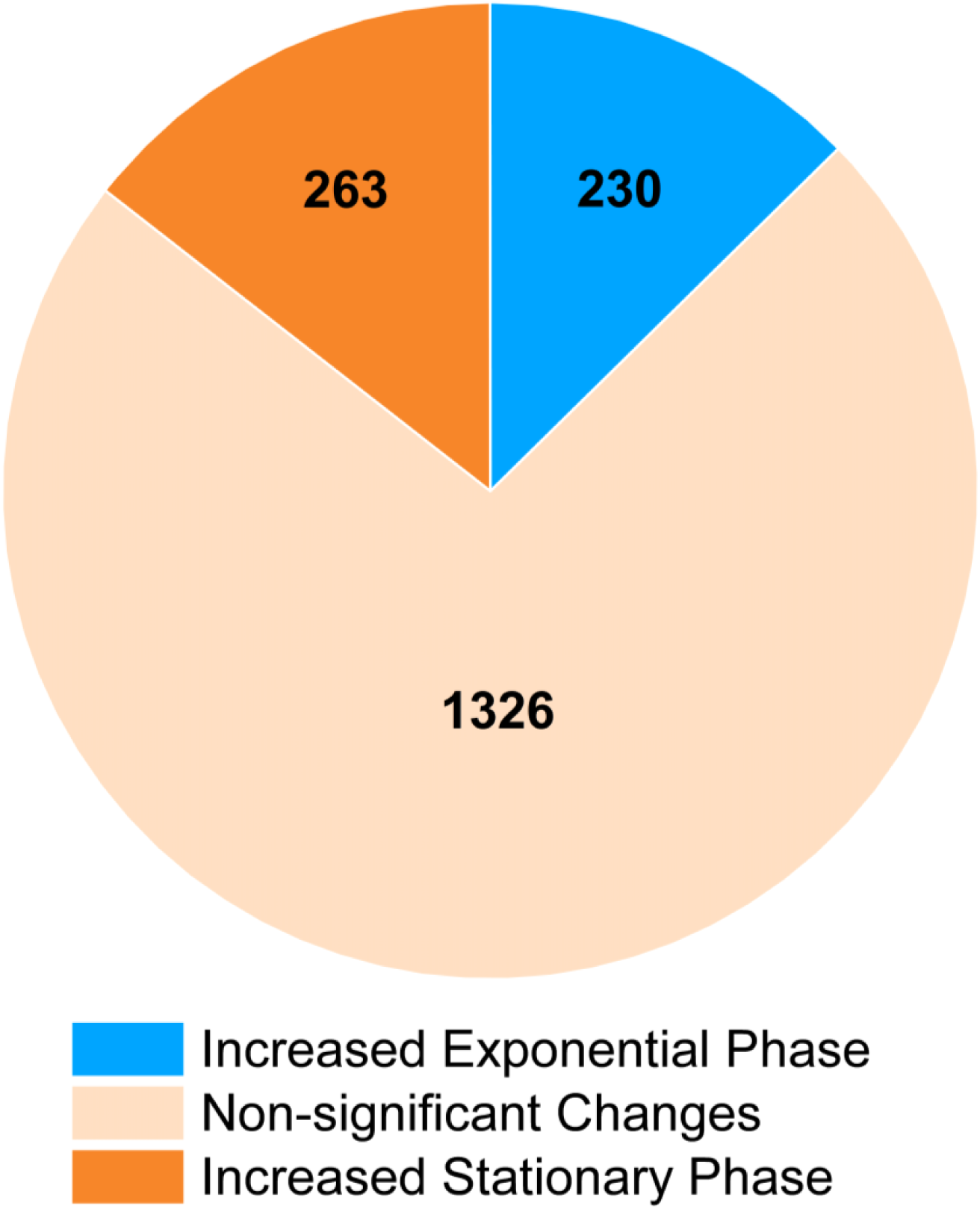
Differential protein abundance in BLR (DE3) *E. coli* cells across growth phases. Of the 1,819 proteins identified, 230 exhibited increased abundance in the exponential phase, while 263 exhibited increased abundance in the stationary phase. The remaining 1,326 proteins displayed no significant change in abundance between the two phases.

In the present study, 230 proteins exhibited biologically and statistically significant increases in abundance during the exponential phase compared to the stationary phase, while 263 showed significant decreases (Figure 1). Protein levels associated with translation, ribosome biogenesis, and gene expression rose during exponential growth to support rapid cell division (Figure 2, Table S1). Conversely, in the nutrient-limited stationary phase, proteins involved in the tricarboxylic acid (TCA) cycle—which enables energy production from alternative carbon sources and provides biosynthetic precursors—were more abundant (Table S2) ^17–19^. Likewise, enzymes involved in small-molecule synthesis that feed into the TCA cycle also increased relative to the exponential phase (Table S2).

**Figure 2.**
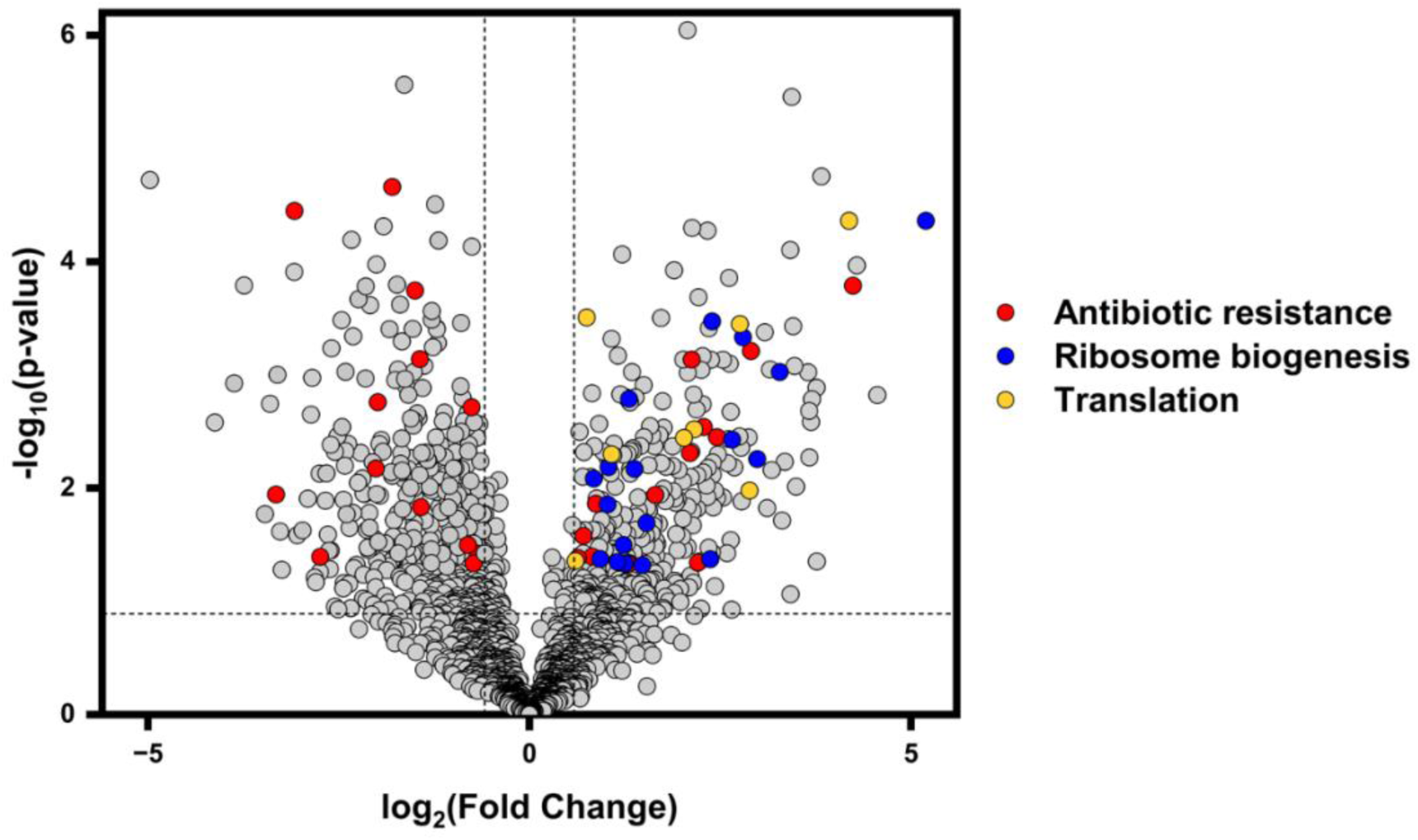
Growth phase-dependent regulation of translation, ribosome biogenesis and antibiotic resistance. Proteins associated with ribosome biogenesis and translation are reduced during the stationary phase, while distinct proteins linked to antibiotic resistance are elevated in both phases. Fold change was calculated as the ratio of protein abundance in the exponential phase to that in the stationary phase. Dotted lines indicate thresholds for biological and statistically significance: a p-value of 0.05 and log₂ (fold change) of ± 0.585. Legend: Proteins involved in antibiotic resistance are shown in red, ribosome biogenesis in blue, and translation in orange.

### Ribosomal maturation factors DeaD, DbpA, RlhE, and RimP are more abundant during the exponential phase than during the stationary phase

The ribosome (70S) in *E. coli* consists of the small (30S) and large (50S) subunits. The small subunit possesses one ribosomal RNA (rRNA) molecule, 16S, and 21 ribosomal proteins, while the large subunit possesses two rRNA molecules, 5S and 23S, and 33 ribosomal proteins ^20^. In bacteria, dozens of proteins transiently interact with ribosomal intermediates and facilitate their maturation ^21–23^. One class of maturation factors that facilitates ribosome assembly in most known organisms is the DEAD-box RNA helicases ^24–28^. In *E. coli*, there are five DEAD-box RNA helicases—DeaD, DbpA, RlhB, RlhE, and SrmB—of which four (DeaD, DbpA, RlhE, and SrmB) are involved in large subunit ribosome assembly; DeaD is also implicated in small subunit assembly ^29–33^. Our data reveal that during the exponential growth phase, the abundance of the DEAD-box RNA helicases DeaD, DbpA, and RlhE increases relative to the stationary phase (Figure 3A).

**Figure 3.**
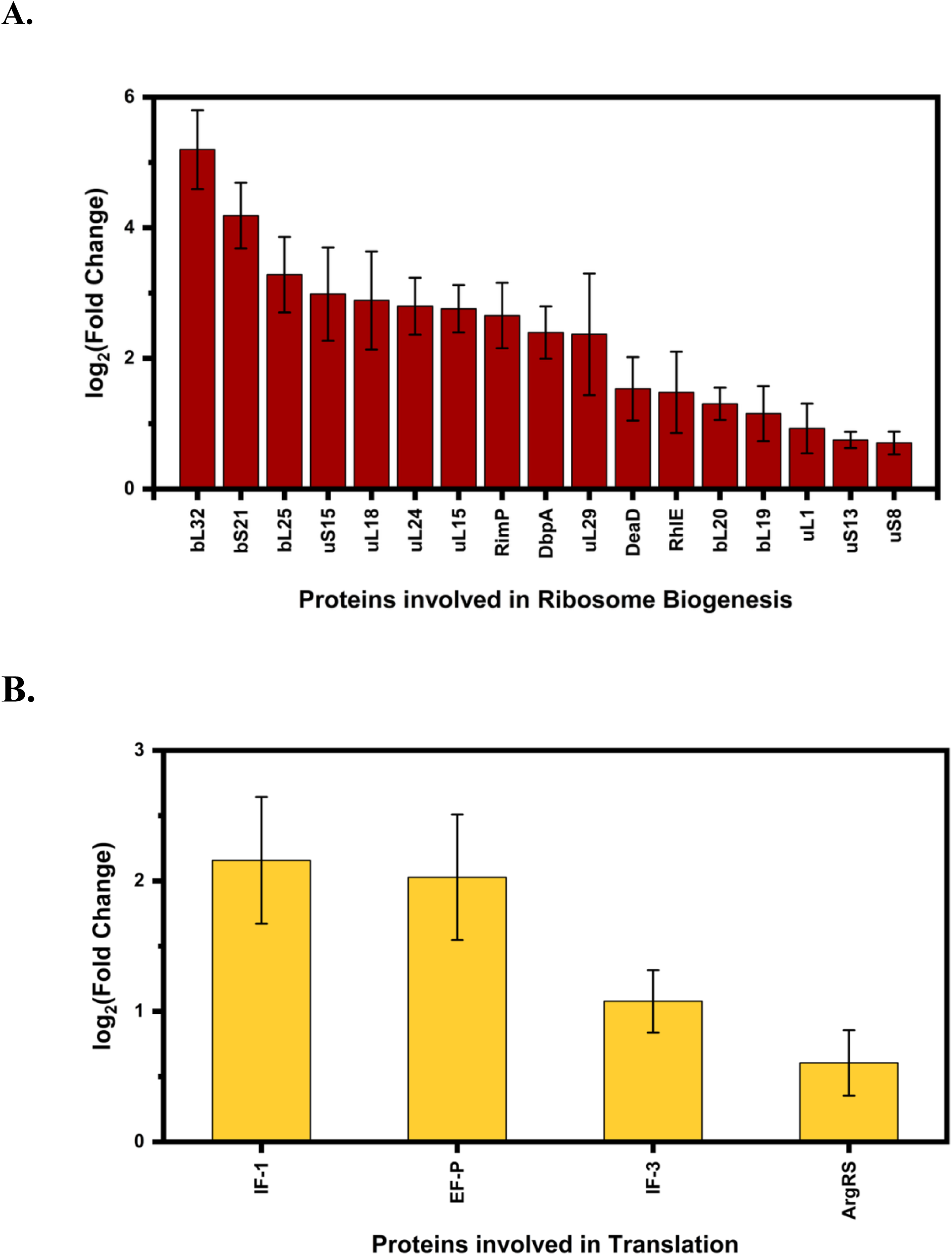
Ribosomal proteins from both the small and large subunits, DEAD-box RNA helicases, and proteins involved in translation modulation are elevated during the exponential phase relative to the stationary phase. A) Ribosomal proteins, DEAD-box RNA helicases, and enzymes involved in rRNA and tRNA modification are elevated during the exponential phase relative to the stationary phase. B) Proteins involved in translation initiation and elongation, along with arginyl-tRNA synthetase (ArgRS), are also elevated during the exponential phase relative to the stationary phase. For both panels A and B, bar graphs represent the log_2_(fold change), and error bars indicate the standard error of the log_2_(fold change). Log_2_(fold change) was calculated as the base-2 logarithm of the ratio of protein abundance in the exponential phase to that in the stationary phase. Proteins were considered differentially abundant if they had a p-value < 0.05 and an absolute log_2_(fold change) ≥ 0.585.

Previous studies have shown that the number of mature ribosomes increases during the fast-growing exponential phase compared to the stationary phase ^34, 35^. Therefore, the increased abundance of DeaD, DbpA, and RlhE likely supports the elevated rate of ribosome production during the exponential growth phase in *E. coli* BLR (DE3) ^35–39^.

In the BW25113 *E. coli* strain, a K-12 strain grown in LB medium, the abundance of the DEAD-box RNA helicases DeaD, RlhE, and SrmB increases during the exponential phase compared to the 24-hour stationary phase ^40^. The DbpA protein was not detected in the BW25113 study ^40^. In our study, similar to the BW25113 strain, the abundance of the DeaD and RlhE proteins increases during the exponential phase (Figure 3A) ^40^. However, unlike the BW25113 study, we observed no change in SrmB abundance between the exponential and stationary phases (Table S3) ^40^.

Our analysis shows that RimP, another protein that facilitates ribosome assembly, is more abundant during the exponential phase compared to the stationary phase (Figure 3A) ^41^. RimP promotes faster binding of small subunit proteins uS9 and uS19 to 30S intermediates, while inhibiting the binding of uS12 and uS13 ^42^. In *E. coli*, deletion of the *rimP* gene results in a high-temperature, slow-growth phenotype and the accumulation of small subunit intermediates ^41^. Our findings regarding RimP protein abundance are consistent with observations in BW25113 cells, where RimP levels also increase during the exponential phase compared to the stationary phase ^40^.

### Exponential phase enrichment of specific ribosomal and translational machinery components

Thirteen large and small ribosomal subunit proteins show increased abundance in the exponential phase compared to the stationary phase (Figure 3A, Table S4). In *E. coli*, several large and small subunit proteins possess RNA and/or protein chaperone activity in addition to their structural roles. Of the thirteen ribosomal proteins more abundant during the exponential phase, uL1 and uL24 exhibit RNA chaperone activity, while uL15, uL18, and bL19 exhibit both RNA and protein chaperone activities ^43, 44^. Therefore, similar to the ribosome maturation factors DeaD, DbpA, RlhE, and RimP, the large subunit proteins uL1, uL15, uL18, bL19, and uL24 likely also facilitate the extensive folding of rRNA and ribosomal proteins necessary for ribosome assembly and protein synthesis during the exponential phase—processes essential for supporting the rapid cell growth and division characteristic of this phase ^35–39^.

Lastly, to support the elevated translational activity characteristic of the exponential phase, initiation factors IF-1 and IF-3, elongation factor EF-P, and ArgRS are more abundant during the exponential phase than in the stationary phase (Figure 3B) ^38, 39^. Similarly, in BW25113 cells, levels of IF-1 and ArgRS also increased during the exponential phase compared to the stationary phase ^40^. In contrast, IF-3 levels remained unchanged between the two phases in BW25113 cells, and EF-P was not detected in that study ^40^.

### RlmN, RsmI, RluB, RlmG, and TsaC—RNA post-transcriptional modifying enzymes—are enriched during the exponential phase

Five enzymes—RlmN, RsmI, RluB, RlmG, and TsaC—implicated in the incorporation of rRNA and tRNA modifications are more abundant in the exponential phase relative to the stationary phase in *E. coli* BLR (DE3) cells (Figure 4) ^45–52^. Three of these RNA-modifying enzymes—RlmN, RluB, and RlmG—act on the 23S rRNA of the large ribosomal subunit ^45, 46, 48, 50, 51^. The RNA-modifying enzyme RsmI acts on the 16S rRNA of the small ribosomal subunit, while TsaC is an essential *E. coli* protein involved in both tRNA modification and small ribosomal subunit assembly ^47, 49, 53, 54^.

**Figure 4.**
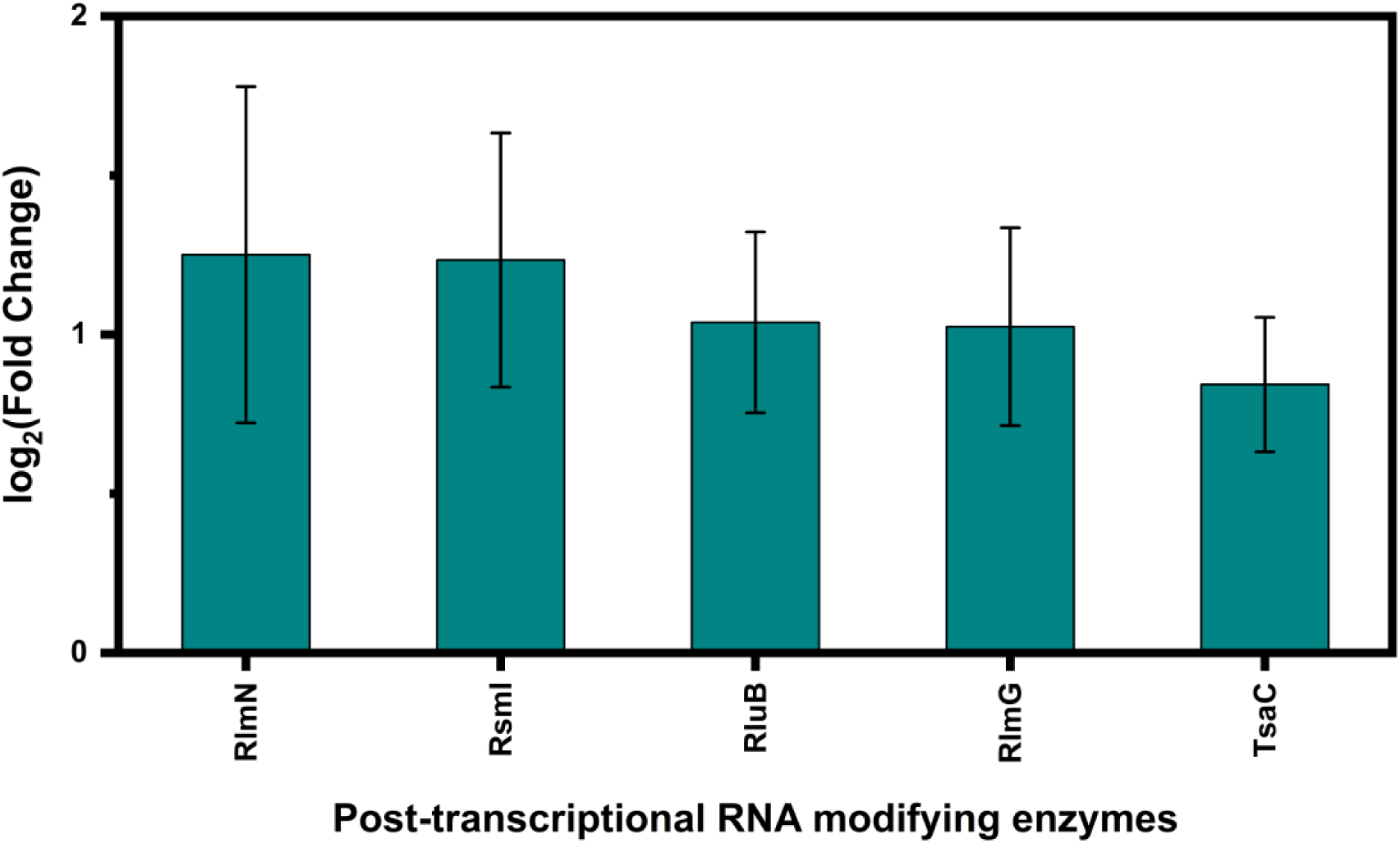
Five RNA post-transcriptional modifying enzymes are enriched during the exponential phase compared to the stationary phase. These enzymes catalyze nucleotide modifications within small and large subunit rRNAs and tRNAs. Bar graphs represent the log_2_(fold change) for each enzyme, and the lines indicate the standard error of the log_2_(fold change). Log_2_(fold change) was calculated as the base-2 logarithm of the ratio of protein abundance in the exponential phase to that in the stationary phase. Proteins were considered differentially abundant if they had a p-value < 0.05 and an absolute log_2_(fold change) ≥ 0.585.

RlmN catalyzes the incorporation of the m^2^A modification at position 2507 of the 23S rRNA^46, 50, 51^. This m^2^A2507 modification—along with the 7-methylguanosine at position 2073 and 2-methylguanosine at position 2447 introduced by RlmKL, the 2′-O-methylcytidine (C_m_) at position 2505 introduced by RlmM, and the pseudouridine at position 2461 introduced by RluE, all located within the peptidyl transferase center (PTC) of 23S rRNA—facilitate maturation of the 50S large ribosomal subunit and increase the translation elongation rate ^45, 46, 55–57^. In DH5α cells, a K-12 strain of *E.coli*, the level of m^2^A2507 is elevated during the exponential growth phase in cells cultured in LB medium, relative to those subjected to metabolic or cold stress ^58^. Similarly, in BW25113 cells, the abundance of RlmN increases during the exponential phase compared to the stationary phase. Thus, during the exponential phase—characterized by enhanced ribosome biogenesis and elevated translation rates—the m^2^A2507 modification, which supports these processes, also increases across multiple *E. coli* strains ^35–39, 56^.

RluB catalyzes the isomerization of U 2609 in 23S rRNA to pseudouridine (Ψ), a modification located within PTC ^52^. The role of RluB in large subunit maturation, as well as the function or significance of the Ψ 2609 modification in ribosome maturation, stability, or translation, remains unclear ^56, 59^. Interestingly, both our data and previous results from BW25113 cells show that RluB is enriched during the exponential phase (Figure 4) ^40^. Although the biological functions of RluB and the Ψ 2609 modification remain unknown, RluB’s selective enrichment during the exponential phase suggests that RluB and/or Ψ 2609 plays a role in optimizing ribosome structure and/or function during rapid cellular proliferation—alternatively, may confer a disadvantage during the stationary phase. Future studies are needed to elucidate how RluB-mediated pseudouridylation modulates ribosome function and structure across different physiological states.

The m²G1837 modification, catalyzed by the RlmG enzyme, has been shown to influence the stability of the *E. coli* 70S ribosome by strengthening interactions between the 50S and 30S subunits ^60, 61^. Similar to the results shown here, the abundance of RlmG is reduced during the stationary phase of BW25113 cells compared to the exponential phase (Figure 4) ^40^. Furthermore, the extent of m²G1837 modification decreases in DH5α cells under cold and metabolic stress conditions ^58^. Collectively, our data and previous studies indicate that under various stress conditions, 70S ribosomes may be less stable and tend to dissociate into 30S and 50S subunits. These free subunits are more vulnerable to degradation, a process that may contribute to energy production during the stationary phase—a growth stage characterized by nutrient limitation ^62^. Additionally, during the stationary phase, a smaller fraction of ribosomes is engaged in active translation ^38, 39^. Thus, the reduced stability of 70S ribosomes during this phase may further increases the pool of inactive 30S and 50S subunits.

RsmI is an enzyme that catalyzes the 2′-O-methylation of the C1402 residue in 16S rRNA ^49^. Another enzyme, RsmH, modifies the same nucleotide by methylating the N4 position of C1402 (m⁴C) ^49^. In BLR (DE3) cells, we observed that RsmI abundance increases during the exponential phase relative to the stationary phase, whereas RsmH levels remain largely unchanged based on our biological and statistical thresholds (Figure 4). These findings suggest that during the exponential phase, a greater proportion of C1402 residues in BLR (DE3) cells are modified by both 2′-O-methylation and N4-methylation. In contrast, during the stationary phase, many C1402 residues likely carry only the N4-methylation. In BW25113 cells, however, the expression levels of both RsmI and RsmH remain constant between the exponential and stationary phases ^40^. Consequently, the proportion of dimethylated C1402 residues in these cells likely remains stable across growth phases.

The C1402 nucleotide makes direct contact with the mRNA positioned in the ribosomal A site of the ribosome ^49^. Both N4-methylation and 2′-O-methylation at this site contribute to maintaining translation fidelity ^49^. Here, we report that the abundance of the RsmI enzyme— which catalyzes 2′-O-methylation of the C1402 residue in the 16S rRNA—decreases during the stationary phase relative to the exponential phase (Figure 4). This finding suggests that, in BLR (DE3) cells, translational accuracy is reduced under stationary-phase conditions. Accordingly, during this growth phase, BLR (DE3) cells may prioritize survival over fidelity in protein synthesis. A decline in translational accuracy under stress conditions has also been observed in other *E. coli* and other bacteria ^63^.

TsaC catalyzes the formation of L-threonylcarbamoyladenylate (t^6^A) ^47^. Subsequently, the TsaB, TsaD, and TsaE enzymes collaboratively incorporate t^6^A at position 37 in tRNA ^47, 64^. This tRNA modification enhances translation fidelity by stabilizing codon–anticodon interactions ^47, 65^. In addition to its role in tRNA modification, the TsaC protein is involved in the maturation of the *E. coli* small ribosomal subunit ^53^. Our data show an increased abundance of the TsaC enzyme in the exponential phase compared to the stationary phase. However, no change was observed in the abundance of TsaB and TsaD, and the TsaE protein was not detected in either growth phase (Table S3). The elevated abundance of TsaC in the exponential phase, without a corresponding increase in TsaD and TsaB, suggests that TsaC may play a role in 30S small subunit assembly or in other, yet-uncharacterized, cellular functions that do not require the activity of the TsaD and TsaB proteins ^53^.

### In *E. coli* BLR (DE3) cells, fourteen proteins associated with antibiotic resistance are more abundant in the exponential phase, while a different set of thirteen proteins is elevated in the stationary phase

The abundances of 27 proteins associated with antibiotic resistance differ between the exponential and stationary phases of *E. coli* BLR (DE3) growth (Figure 5, Table 1). These proteins are involved in wide rage antibiotic resistance mechanisms and cellular metabolic pathways (Table S5-S8). Notably, several proteins exhibit similar abundance patterns between the two growth phases in both BLR (DE3) and BW25113 strains, whereas others either remain unchanged in BW25113 or display opposite trends in relative abundance ^40^.

**Figure 5.**
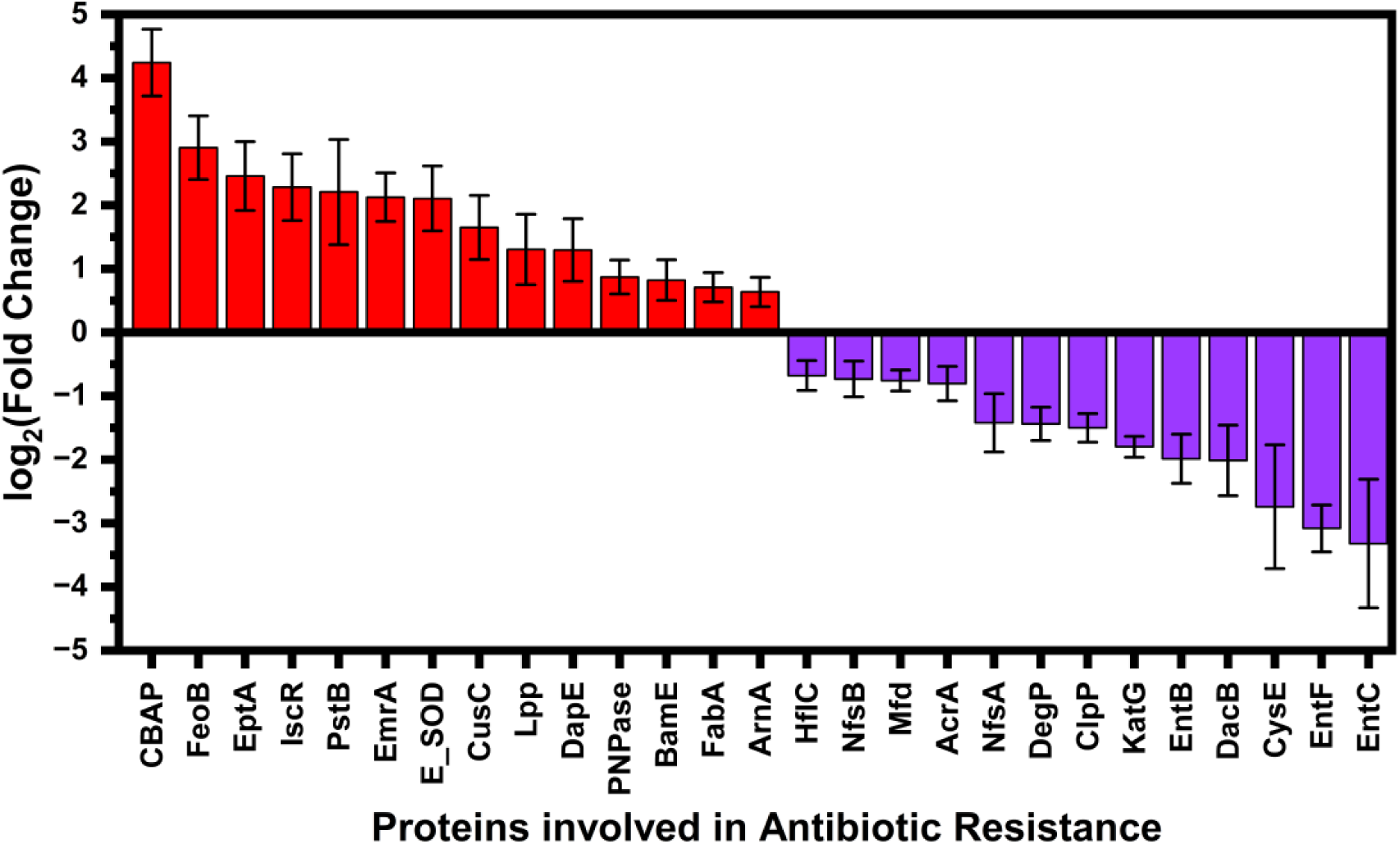
Protein expression profiles reveal growth phase–specific antibiotic resistance mechanisms. Fourteen proteins, indicated in red, are upregulated in the exponential phase relative to the stationary phase. Thirteen proteins, indicated in purple, are upregulated in the stationary phase relative to the exponential phase. Bar graphs represent the log_2_(fold change) for each enzyme and the lines indicate the standard error of the log_2_(fold change). Log_2_(fold change) was calculated as the base-2 logarithm of the ratio of protein abundance in the exponential phase to that in the stationary phase. Proteins were considered differentially abundant if they had a p-value < 0.05 and an absolute log_2_ (fold change) ≥ 0.585.

**Table 1.**
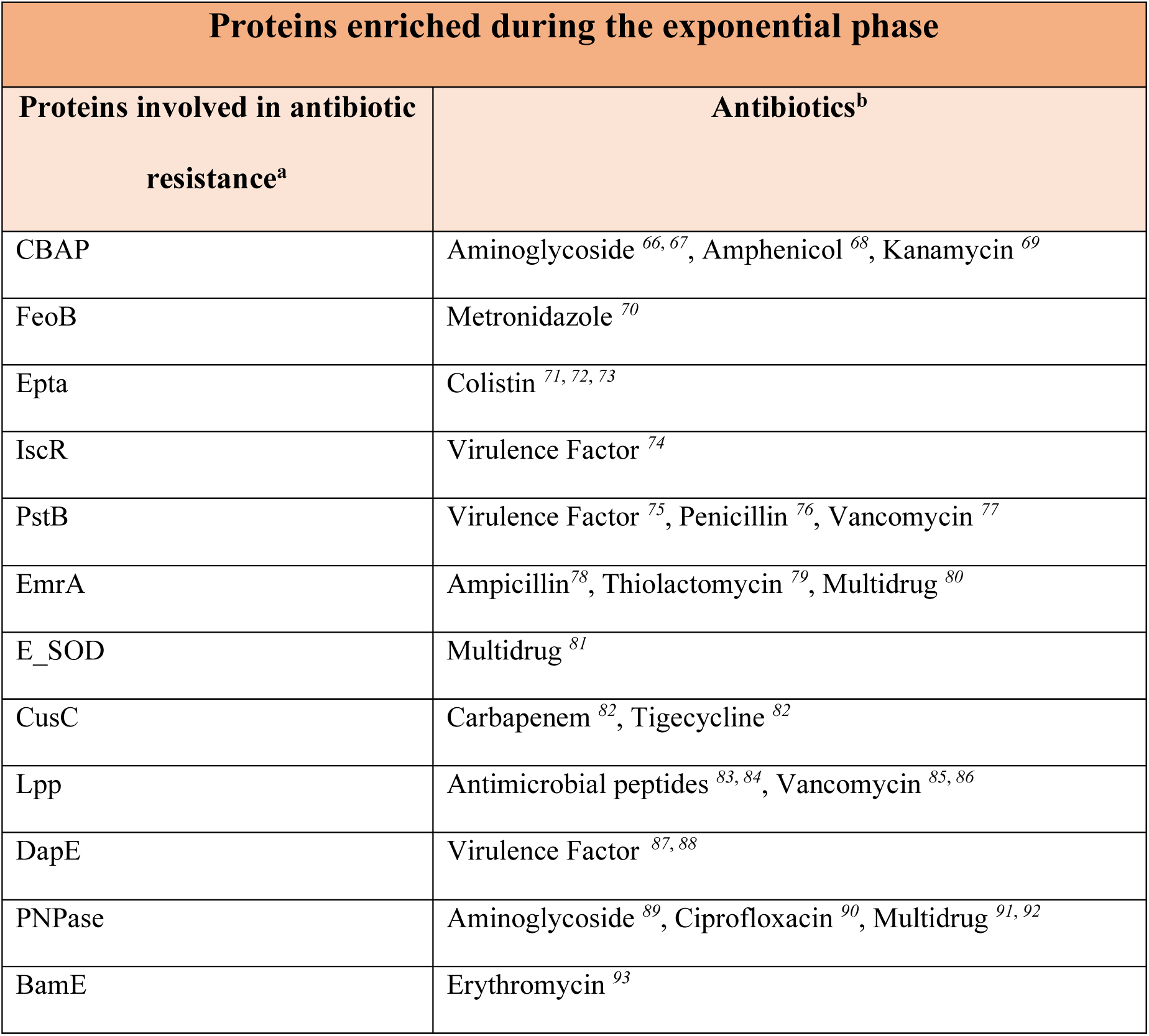
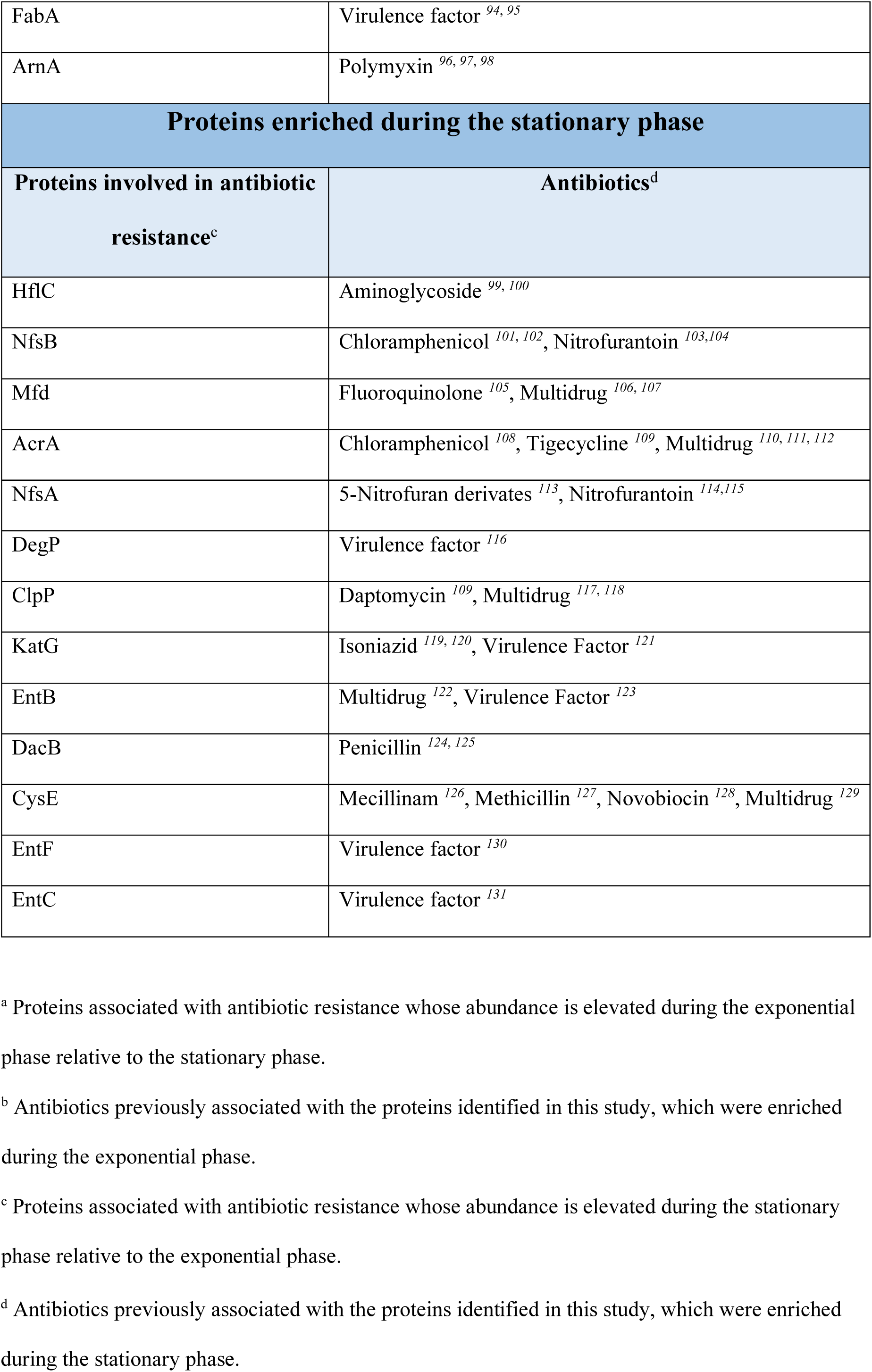
Unique proteins associated with single- and multidrug resistance are enriched during the exponential or stationary growth phase in *E. coli* BLR (DE3).

The abundance of seven proteins — CBAP, FeoB, ArnA, EmrA, IscR, PstB, and DapE — is higher in the exponential phase of both BLR (DE3) and BW25113 cells relative to the stationary phase (Figure 5, Table 1) ^40^. On the other hand, the abundance of six proteins — E_SOD, CusC, Lpp, BamE, FabA, and PNPase — increases in the exponential phase of BLR (DE3) cells relative to the stationary phase, but the abundance of these proteins exhibits the opposite trend in the BW25113 cells, or remains unchanged between the two phases (Figure 5, Table 1) ^40^. The abundance of the EptA protein increases in the exponential phase of BLR (DE3) cells, while this protein is not detected in the BW25113 cells (Figure 5, Table 1) ^40^.

In the stationary phase, the abundance of three proteins, EntB, EntC, and EntF, increases relative to the exponential phase for both BLR (DE3) and BW25113 cells (Figure 5, Table 1) ^40^. In contrast, the abundance of nine proteins, HflC, Mfd, AcrA, NsfA, DegP, ClpP, KatG, DacB and CysE increases in the stationary phase of BLR (DE3) cells relative to the exponential phase while these abundances remain unchanged or show the opposite trend in the BW25113 cells. Finally, the abundance of the NsfB protein increases in the stationary phase in of the BLR (DE3) cells whereas its abundance remains undetermined in BW25113 cells ^40^.

In summary, the data collected in this study, combined with previous results, demonstrate that the two different *E. coli* strains—BLR (DE3), a B strain, and BW25113, a K-12 strain — possess a number of shared and many unique, antibiotic resistance mechanisms during the exponential and stationary growth phases ^40^.

### Conclusion

Our results are consistent with previous findings in *E. coli*, which show that processes essential for rapid cell growth—such as gene expression, ribosome assembly, and translation—are elevated in the exponential phase relative to the stationary phase (Figure 1, Table S1) ^35–39^. In contrast, the TCA cycle, which is required for energy production and biosynthesis processes when glucose is limited, is more active in the stationary phase than in the exponential phase (Table S2) ^17–19^.

Importantly, we conducted a systematic and comprehensive analysis of differences in antibiotic resistance mechanisms between the exponential and stationary phases. Our results demonstrate that distinct resistance pathways are activated in each phase of BLR (DE3), a nonpathogenic *E. coli* B strain. This insight may inform the development of phase-specific antibiotics targeting pathogenic *E. coli* and other bacteria (Figure 5, Table 1).

Lastly, we observed distinct differences in the proteomic profiles associated with antibiotic resistance between the nonpathogenic *E. coli* B strain characterized in this study and a previously investigated nonpathogenic *E. coli* K-12 strain ^40^. Future large-scale, quantitative proteomic analyses comparing pathogenic *E. coli* strains with nonpathogenic *E. coli* from the human microbiome could uncover both conserved and strain-specific resistance pathways. This knowledge may inform the development of narrow-spectrum antibiotic treatments that selectively target pathogenic *E. coli* while preserving commensal strains, thereby minimizing microbiome disruption and reducing the potential for resistance development.

## Data Availability

The mass spectrometry proteomics data and associated metadata, experimental details, and sample-to-data files have been deposited to the ProteomeXchange Consortium via the PRIDE partner repository with the dataset identifier XXXXXX ^132, 133^.

## Associated Content Supporting Information

Biological pathways elevated in the exponential phase of *E. coli* BLR (DE3) cells relative to the stationary phase. Biological pathways elevated in the stationary phase of *E. coli* BLR (DE3) cells relative to the exponential phase. Log₂ fold change, p-values, and the phase-specific abundance of TsaB, TsaD, RsmH, and SrmB proteins in the exponential and stationary growth phases of *E. coli*. Relative abundance and p-values of ribosomal structural proteins. Cellular functions of proteins whose abundance uniquely increases during the exponential phase of *E. coli* growth and that are involved in antibiotic resistance. Cellular functions of proteins whose abundance uniquely increases during the stationary phase of *E. coli* growth and that are involved in antibiotic resistance. Biological processes associated with proteins involved in antibiotic resistance that increase in abundance during the exponential phase. Biological processes associated with proteins involved in antibiotic resistance that increase in abundance during the stationary phase.

## Acknowledgment

EK is grateful to Luis Gracia Mazuca for the discussions concerning the RNA modifying enzymes.

---

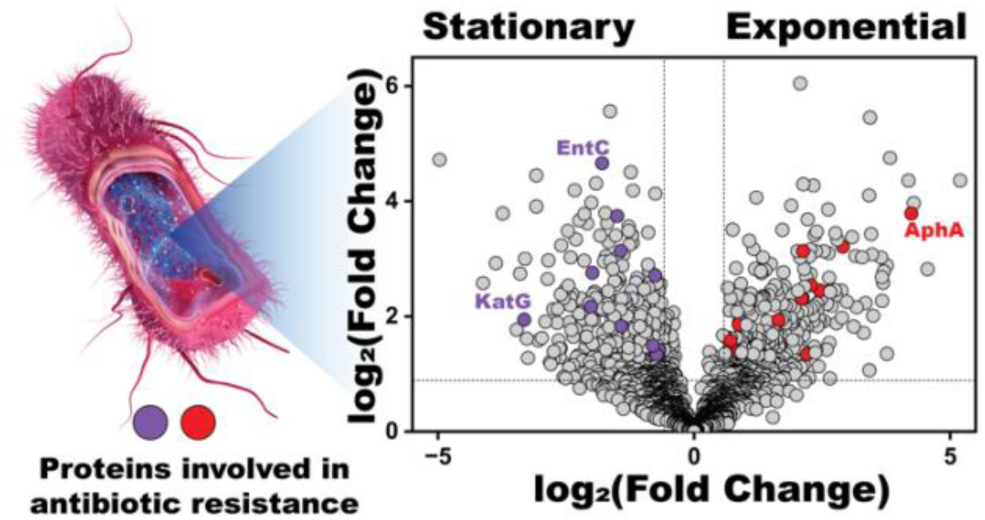
FOR TABLE OF CONTENT USE ONLY.

