## Supporting Information for "Phase-Specific Antibiotic Resistance Mechanisms in an *Escherichia coli* B Strain"

**Table of Contents**:

**Table S1**. Biological pathways elevated in the exponential phase of *E. coli* BLR (DE3) cells relative to the stationary phase.

**Table S2**. Biological pathways elevated in the stationary phase of *E. coli* BLR (DE3) cells relative to the exponential phase.

**Table S3** $Log_2$(fold change), p-values, and phase-specific abundance of TsaB, TsaD, RsmH, and SrmB proteins in exponential and stationary growth phases of *E. coli.*

**Table S4.** Relative abundance and p-values of ribosomal structural proteins.

**Table S5**. Cellular functions of proteins whose abundance uniquely increases during the exponential phase of *E. coli* growth that are involved in antibiotic resistance.

**Table S6**. Cellular functions of proteins whose abundance uniquely increases during the stationary phase of *E. coli* growth that are involved in antibiotic resistance.

**Table S7**. Biological processes associated with proteins involved in antibiotic resistance that increase in abundance during the exponential phase.

**Table S8**. Biological processes associated with proteins involved in antibiotic resistance that increase in abundance during the stationary phase.

**Table S1**. Biological pathways elevated in the exponential phase of *E. coli* BLR (DE3) cells relative to the stationary phase.

| **Pathways increased in exponential phase** | | | | | |
|---|---|---|---|---|---|
| **Biological Process**[a] | **GO-term**[b] | **Count in network**[c] | **Strength**[d] | **Signal**[e] | **False Discovery Rate**[f] |
| Ribosome assembly | GO:0042255 | 14 of 55 | 0.67 | 0.51 | 0.0057 |
| Ribonucleoprotein complex assembly | GO:0022618 | 13 of 51 | 0.67 | 0.49 | 0.0069 |
| Ribosome biogenesis | GO:0042254 | 20 of 104 | 0.54 | 0.48 | 0.0057 |
| Post-transcriptional regulation of gene expression | GO:0010608 | 15 of 71 | 0.59 | 0.46 | 0.0083 |
| Cytoplasmic translation | GO:0002181 | 13 of 57 | 0.62 | 0.44 | 0.0117 |
| Ribosomal large subunit assembly | GO:0000027 | 9 of 29 | 0.75 | 0.42 | 0.0179 |
| Cellular component biogenesis | GO:0044085 | 39 of 32 | 0.34 | 0.41 | 0.0057 |
| Cellular component organization or biogenesis | GO:0071840 | 50 of 459 | 0.3 | 0.4 | 0.0057 |
| Cellular component organization | GO:0016043 | 44 of 398 | 0.3 | 0.4 | 0.0057 |
| Non-membrane-bounded organelle assembly | GO:0140694 | 15 of 79 | 0.54 | 0.4 | 0.0169 |
| Gene expression | GO:0010467 | 39 of 354 | 0.3 | 0.37 | 0.0110 |
| Nitrogen compound metabolic process | GO:0006807 | 116 of 1464 | 0.16 | 0.36 | 0.0053 |
| Cellular macromolecule metabolic process | GO:0044260 | 59 of 625 | 0.24 | 0.36 | 0.0083 |
| Cellular macromolecule biosynthetic process | GO:0034645 | 34 of 294 | 0.32 | 0.36 | 0.0131 |
| Cellular component assembly | GO:0022607 | 28 of 227 | 0.35 | 0.35 | 0.0179 |
| Organelle organization | GO:0006996 | 20 of 139 | 0.42 | 0.35 | 0.0239 |
| Translation | GO:0006412 | 18 of 120 | 0.44 | 0.33 | 0.0308 |
| Cellular nitrogen compound metabolic process | GO:0034641 | 80 of 986 | 0.17 | 0.31 | 0.0179 |

| | | | | | |
|---|---|---|---|---|---|
| Primary metabolic process | GO:0044238 | 120 of 1653 | 0.12 | 0.3 | 0.0169 |
| Cellular metabolic process | GO:0044237 | 133 of 1910 | 0.1 | 0.28 | 0.0234 |
| Macromolecule metabolic process | GO:0043170 | 80 of 1012 | 0.16 | 0.27 | 0.0332 |
| Organic substance metabolic process | GO:0071704 | 133 of 1934 | 0.1 | 0.25 | 0.0376 |
| **Molecular Function**[a] | | | | | |
| **Biological Process** | **GO-term** | **Count in network**[1] | **Strength**[2] | **Signal**[3] | **False Discovery Rate**[4] |
| Structural molecule activity | GO:0005198 | 17 of 72 | 0.63 | 0.55 | 0.0027 |
| rRNA binding | GO:0019843 | 15 of 60 | 0.66 | 0.53 | 0.0040 |
| RNA binding | GO:0003723 | 29 of 184 | 0.46 | 0.5 | 0.0027 |
| Structural constituent of ribosome | GO:0003735 | 13 of 58 | 0.61 | 0.37 | 0.0282 |
| Binding | GO:0005488 | 151 of 2169 | 0.1 | 0.31 | 0.0114 |
| **Cellular Component**[a] | | | | | |
| **Biological Process** | **GO-term** | **Count in network**[1] | **Strength**[2] | **Signal**[3] | **False Discovery Rate**[4] |
| Ribonucleoprotein complex | GO:1990904 | 14 of 61 | 0.62 | 0.6 | 0.0014 |
| Periplasmic space | GO:0042597 | 33 of 250 | 0.38 | 0.54 | 0.00058 |
| Outer membrane-bounded periplasmic space | GO:0030288 | 27 of 188 | 0.42 | 0.54 | 0.00090 |
| Cytosol | GO:0005829 | 96 of 1066 | 0.22 | 0.53 | 0.0000383 |
| Ribosomal subunit | GO:0044391 | 13 of 59 | 0.6 | 0.53 | 0.0034 |
| Cytosolic ribosome | GO:0022626 | 13 of 60 | 0.6 | 0.53 | 0.0034 |
| Cytoplasm | GO:0005737 | 118 of 1443 | 0.17 | 0.5 | 0.0000383 |
| Cytosolic large ribosomal subunit | GO:0022625 | 9 of 34 | 0.68 | 0.48 | 0.0078 |
| Cell envelope | GO:0030313 | 35 of 311 | 0.31 | 0.43 | 0.0034 |
| Non-membrane-bounded organelle | GO:0043228 | 21 of 161 | 0.38 | 0.4 | 0.0097 |
| Cellular anatomical entity | GO:0110165 | 185 of 2904 | 0.07 | 0.34 | 0.0034 |

[a] The data presented in this table were obtained from STRING following the upload of all proteins that exhibited a $\log_2$(fold change) > 0.585 and a p-value < 0.05 (Figure 1) [1]. The fold change was calculated as the ratio of protein abundance in the exponential phase to that in the stationary phase

[b] GO-term is the Gene Ontology accession number [2, 3].

[c] The first number represents the proteins that are upregulated during the exponential phase and annotated with a specific GO term. The second number refers to the total number of *E. coli* K-12 proteins associated with that GO term [1].

[d] Strength is calculated as $\log_{10}$ (observed/expected), where "observed" denotes the number of proteins within the STRING-derived network that are annotated with a specific Gene Ontology (GO) term, and "expected" refers to the number of such proteins anticipated in a randomly generated network of equivalent size. The expected value is based on the overall frequency of the annotation in the complete *E. coli* proteome. A higher strength score indicates that the annotation occurs more frequently in the STRING-derived network than would be expected by chance, suggesting functional enrichment [1].

[e] Signal is computed as the average of the strength score and the negative logarithm (base 10) of the false discovery rate (FDR). A high signal indicates both a strong enrichment (high strength) and a low FDR, implying a low expected proportion of false positives among all statistically significant annotations [1].

[f] The false discovery rate estimates the expected proportion of false positives among all statistically significant annotations [1].

**Table S2**. Biological pathways elevated in the stationary phase of *E. coli* BLR (DE3) cells relative to the exponential phase.

| **Pathways increased in stationary phase** | | | | | |
|---|---|---|---|---|---|
| **Biological Process** | **GO-term** | **Count in network** | **Strength** | **Signal** | **False Discovery Rate** |
| Tricarboxylic acid cycle | GO:0006099 | 12 of 29 | 0.82 | 0.54 | 0.0058 |
| Aerobic respiration | GO:0009060 | 15 of 59 | 0.61 | 0.46 | 0.0083 |
| Small molecule biosynthetic process | GO:0044283 | 42 of 309 | 0.34 | 0.42 | 0.0058 |
| Carboxylic acid metabolic process | GO:0019752 | 60 of 527 | 0.26 | 0.39 | 0.0058 |
| Oxoacid metabolic process | GO:0043436 | 61 of 556 | 0.24 | 0.37 | 0.0078 |
| Small molecule metabolic process | GO:0044281 | 86 of 892 | 0.19 | 0.35 | 0.0083 |
| Cellular metabolic process | GO:0044237 | 156 of 1910 | 0.12 | 0.34 | 0.0058 |
| Metabolic process | GO:0008152 | 172 of 2134 | 0.11 | 0.34 | 0.0058 |
| Organic substance metabolic process | GO:0071704 | 152 of 1934 | 0.1 | 0.26 | 0.0366 |
| **Molecular Function** | | | | | |
| **Biological Process** | **GO-term** | **Count in network[1]** | **Strength[2]** | **Signal[3]** | **False Discovery Rate[4]** |
| Small molecule binding | GO:0036094 | 107 of 834 | 0.31 | 0.82 | 1.50e-10 |
| Anion binding | GO:0043168 | 94 of 732 | 0.31 | 0.78 | 6.56e-09 |
| Nucleotide binding | GO:0000166 | 93 of 720 | 0.31 | 0.78 | 6.56e-09 |
| Heterocyclic compound binding | GO:1901363 | 142 of 1439 | 0.2 | 0.63 | 4.65e-08 |
| Organic cyclic compound binding | GO:0097159 | 142 of 1439 | 0.2 | 0.63 | 4.65e-08 |
| Carbohydrate derivative binding | GO:0097367 | 71 of 579 | 0.29 | 0.62 | 9.90e-06 |
| Protein binding | GO:0005515 | 64 of 505 | 0.31 | 0.62 | 1.75e-05 |
| Ribonucleotide binding | GO:0032553 | 68 of 553 | 0.29 | 0.61 | 1.75e-05 |
| Catalytic activity | GO:0003824 | 176 of 1963 | 0.16 | 0.59 | 3.14e-08 |
| Binding | GO:0005488 | 189 of 2169 | 0.14 | 0.58 | 1.81e-08 |
| Ion binding | GO:0043167 | 130 of 1338 | 0.19 | 0.57 | 1.71e-06 |

| | | | | | |
|---|---|---|---|---|---|
| Identical protein binding | GO:0042802 | 56 of 441 | 0.31 | 0.57 | 0.00010 |
| Adenyl ribonucleotide binding | GO:0032559 | 57 of 451 | 0.31 | 0.57 | 0.00010 |
| ATP binding | GO:0005524 | 56 of 445 | 0.3 | 0.56 | 0.00012 |
| NAD binding | GO:0051287 | 14 of 55 | 0.61 | 0.51 | 0.0044 |
| Purine ribonucleoside triphosphate binding | GO:0035639 | 57 of 493 | 0.27 | 0.47 | 0.0010 |
| Purine ribonucleotide binding | GO:0032555 | 59 of 517 | 0.26 | 0.47 | 0.0010 |
| Oxidoreductase activity, acting on the CH-OH group of donors, NAD or NADP as acceptor | GO:0016616 | 18 of 91 | 0.5 | 0.46 | 0.0062 |
| Oxidoreductase activity | GO:0016491 | 48 of 438 | 0.24 | 0.33 | 0.0177 |
| **Cellular Component** | | | | | |
| **Biological Process** | **GO-term** | **Count in network[1]** | **Strength[2]** | **Signal[3]** | **False Discovery Rate[4]** |
| Cytosol | GO:0005829 | 130 of 1066 | 0.29 | 0.86 | |
| Intracellular anatomical structure | GO:0005622 | 166 of 1487 | 0.25 | 0.83 | |
| Cytoplasm | GO:0005737 | 162 of 1443 | 0.25 | 0.83 | |
| Cellular anatomical entity | GO:0110165 | 212 of 2904 | 0.07 | 0.36 | |

[a] The data presented in this table were obtained from STRING following the upload of all proteins that exhibited a $\log_2$(fold change) < - 0.585 and a p-value < 0.05 (Figure 1) [1]. The fold change was calculated as the ratio of protein abundance in the exponential phase to that in the stationary phase (Figure 1) [1].

[b] GO-term is the Gene Ontology accession number [2, 3].

[c] The first number represents the proteins that are upregulated during the stationary phase and annotated with a specific GO term. The second number refers to the total number of *E. coli* K-12 proteins associated with that GO term [1].

[d] Strength is calculated as explained in Table S1 legend.

[e] Signal is calculated as explained in Table S1 legend.

[f] The false discovery rate estimates the expected proportion of false positives among all statistically significant annotations [1].

**Table S3**. $Log_2$(fold change), p-values, and phase-specific abundance of TsaB, TsaD, RsmH, and SrmB proteins in exponential and stationary growth phases of *E. coli.*

| **Description** | **Gene[a]** | **$Log_2$ FC[b]** | **SEM[c]** | ***p*-value[d]** | **Mean 3 h[e]** | **Mean 22 h[e]** |
|---|---|---|---|---|---|---|
| tRNA threonylcarbamoyladenosine biosynthesis protein TsaB | ***tsaB*** | 1.177 | 0.6797 | 0.134222 | 23.72 | 22.55 |
| tRNA N6-adenosine threonylcarbamoyltransferase | ***tsaD*** | 0.5867 | 0.4395 | 0.224905 | 25.09 | 24.51 |
| Ribosomal RNA small subunit methyltransferase H | ***rsmH*** | -0.2401 | 0.3884 | 0.558947 | 25.21 | 25.45 |
| ATP-dependent RNA helicase SrmB | ***srmB*** | -0.5152 | 0.2579 | 0.110605 | 25.13 | 25.65 |

[a] The name of the gene.

[b] $Log_2FC$ is the base-2 logarithm of the fold-change. The fold-change was calculated as the ratio of protein abundance in the exponential phase to that in the stationary phase.

[c] SEM is the standard error of the $log_2FC$.

[d] The p-value was calculated as explained in the Materials and Methods section of the manuscript. The thresholds for identifying a significant change in protein abundance between the exponential and stationary phases are a $p\text{-value} < 0.05$ and an absolute $log_2$(fold change) $\geq 0.585$**.**

*[e]* The mean values were calculated from protein abundance measured across five replicates. The data were obtained during the exponential phase (3 h) and the stationary phase (22 h). In this table, "h" refers to hours.

**Table S4.** Log2(fold change) and p-values of ribosomal structural proteins.

| Gene (Protein)[a] | $Log_2FC$[b] | *p*-value[c] |
|---|---|---|
| **Small Ribosomal Subunit 30S** | | |
| *rpsA* (bS1) | 1.297 | 0.094025 |
| *rpsB* (uS2) | 0.3029 | 0.571245 |
| *rpsC* (uS3) | 0.7615 | 0.097251 |
| *rpsD* (uS4) | 0.3053 | 0.411026 |
| *rpsE* (uS5) | -0.4535 | 0.196395 |
| *rpsF* (bS6) | -0.5432 | 0.475198 |
| *rpsG* (uS7) | 0.1181 | 0.795367 |
| *rpsH* (uS8) | 0.704 | 0.006399 |
| *rpsI* (uS9) | 0.05746 | 0.891323 |
| *rpsJ* (uS10) | 0.5031 | 0.320432 |
| *rpsK* (uS11) | 0.02624 | 0.986015 |
| *rpsL* (uS12) | 0.6624 | 0.721344 |
| *rpsM* (uS13) | 0.7511 | 0.00031 |
| *rpsN* (uS14) | 0.5344 | 0.467731 |
| *rpsO* (uS15) | 2.984 | 0.005522 |
| *rpsP* (bS16) | -0.8117 | 0.471724 |
| *rpsQ* (uS17) | -1.513 | 0.009966 |
| *rpsR* (bS18) | -1.309 | 0.125283 |
| *rpsS* (uS19) | 0.4328 | 0.661995 |
| *rpsT* (bS20) | -0.5843 | 0.364828 |
| *rpsU* (bS21) | 4.188 | 0.000043 |
| **Large Ribosomal Subunit 50S** | | |
| *rplA* (uL1) | 0.9274 | 0.042273 |
| *rplB* (uL2) | -0.1131 | 0.726299 |
| *rplC* (uL3) | 0.5662 | 0.11742 |
| *rplD* (uL4) | -0.2267 | 0.661021 |
| *rplE* (uL5) | 0.02892 | 0.94639 |
| *rplF* (uL6) | 0.6019 | 0.339576 |
| *rplI* (bL9) | 1.319 | 0.176474 |
| *rplJ* (uL10) | 0.6538 | 0.330838 |
| *rplK* (uL11) | 0.5043 | 0.275507 |
| *rplL* (bL12) | 0.3731 | 0.66252 |
| *rplM* (uL13) | 0.2813 | 0.628971 |
| *rplN* (uL14) | 0.2052 | 0.554677 |
| *rplO* (uL15) | 2.76 | 0.000357 |
| *rplP* (uL16) | -0.4303 | 0.235698 |
| *rplQ* (bL17) | -0.0608 | 0.86579 |
| *rplR* (uL18) | 2.887 | 0.0105 |
| *rplS* (bL19) | 1.154 | 0.0448 |
| *rplT* (bL20) | 1.305 | 0.001632 |
| *rplU* (bL21) | -0.9903 | 0.098794 |
| *rplV* (uL22) | 0.2615 | 0.389387 |
| *rplW* (uL23) | ---- | ---- |
| *rplX* (uL24) | 2.799 | 0.000465 |

| | | |
|---|---|---|
| *rplY* (bL25) | 3.282 | 0.000941 |
| *rpmA* (bL27) | 0.3813 | 0.555182 |
| *rpmB* (bL28) | -1.452 | 0.113618 |
| *rpmC* (uL29) | 2.369 | 0.042377 |
| *rpmD* (uL30) | -1.8 | 0.181018 |
| *rpmF* (bL32) | 5.197 | 0.000043 |
| *rpmG* (bL33) | ---- | ---- |
| *rpmH* (bL34) | ---- | ---- |

[a] The names of the genes encoding the ribosomal structural proteins, with the names of the corresponding proteins shown in parentheses.

[b] $Log_2FC$ is the base-2 logarithm of the fold change, calculated as the ratio of protein abundance in the exponential phase to that in the stationary phase.

[c] The p-value was calculated as described in the Materials and Methods section of the manuscript. Our thresholds for considering a change in protein abundance between the exponential and stationary phases are a p-value $< 0.05$ and an absolute $\log_2FC > 0.585$. Based on these thresholds, no significant difference in protein abundance between the two phases was observed for proteins not highlighted in this table. Proteins highlighted in orange are elevated in abundance in the exponential phase relative to the stationary phase. Proteins highlighted in blue are elevated in abundance in the stationary phase relative to the exponential phase. Proteins highlighted in gray were not detected in this study.

**Table S5**. Cellular functions of proteins whose abundance uniquely increases during the exponential phase of *E. coli* growth that are involved in antibiotic resistance.

| Protein | Function[a] | UniProt[b] |
|---|---|---|
| **CBAP** | Removes phosphate groups from organic monophosphate esters. | P0AE22 |
| **FeoB** | Functions as a transporter in a GTP-driven $Fe^{2+}$ uptake system; probably couples GTP binding to channel opening for iron import. | P33650 |
| **EptA** | Transfers a phosphoethanolamine group to lipid A, this modification is essential for resistance to polymyxin antibiotics. | P30845 |
| **IscR** | Regulates the transcription of multiple genes and operons involved in the biogenesis of Fe-S cluster-containing proteins. | P0AGK8 |
| **PstB** | A component of the PstSACB ABC transport complex, it provides energy for the active uptake of phosphate. | P0AAH0 |
| **EmrA** | Part of the EmrAB-TolC multidrug efflux system, which expels toxic compounds and confers resistance to antibiotics such as CCCP, FCCP, 2,4-dinitrophenol, and nalidixic acid. | P27303 |
| **E_SOD** | Neutralizes harmful ROS within the cell, protecting cellular components from oxidative damage. | P0AGD1 |
| **CusC** | Forms outer membrane pores that enable passive diffusion of metal cations like copper and silver ions | P77211 |
| **Lpp** | Outer membrane lipoprotein that maintains periplasmic space by regulating the distance between inner and outer membranes; adding residues increases periplasm width. | P69776 |
| **DapE** | Catalyzes the cleavage of SDAP, producing intermediates crucial for the synthesis of lysine and *meso*-diaminopimelic acid, which are necessary for peptidoglycan biosynthesis, an essential component of bacterial cell walls. | P0AED7 |
| **PNPase** | Degrades mRNA from the 3′ to 5′ end and also plays roles in tRNA processing and rRNA quality control, especially during nutrient limitation and steady-state growth. | P05055 |
| **BamE** | Member of the beta-barrel assembly machinery complex, which integrates β-barrel proteins into the bacterial outer membrane. | P0A937 |
| **FabA** | Required for the introduction of cis unsaturation into fatty acids | P0A6Q3 |
| **ArnA** | A bifunctional enzyme that modifies UDP-glucuronic acid into forms of L-arabinose derivatives, which are added to lipid A to confer resistance to polymyxin and cationic antimicrobial peptides. | P77398 |

[a] The protein function was obtained from UniProt [4].

[b] The UniProt accession code for each protein [4].

**Table S6**. Cellular functions of proteins whose abundance uniquely increases during the stationary phase of *E. coli* growth that are involved in antibiotic resistance.

| Protein | Function[a] | UniProt[b] |
|---|---|---|
| **HflC** | HflC and HflK, modulates the stability of the bacteriophage lambda cII regulatory protein, thereby influencing the frequency of lysogeny. | P0ABC3 |
| **NfsB** | Functions as a nitroreductase that catalyzes the reduction of a broad range of nitroaromatic substrates using NADH (primarily) and NADPH (to a lesser extent) as electron donors, two electrons are transferred. Capable of reducing nitrofurazone, quinones and the anti-tumor agent CB1954 (5-(aziridin-1-yl)-2,4-dinitrobenzamide). | P38489 |
| **Mfd** | Couples transcription and DNA repair by recognizing RNA polymerase (RNAP) stalled at DNA lesions. Promotes ATP-dependent disassociation of the RNA polymerase complex and facilitates recruitment of the nucleotide excision repair (NER) machinery to the lesion. | P30958 |
| **AcrA** | Component of the AcrA-AcrB-AcrZ-TolC multidrug efflux complex, uses the proton motive force to export substrates. | P0AE06 |
| **NfsA** | Catalyzes the NADPH-dependent reduction of nitroaromatic compounds through a ping-pong bi-bi mechanism. Has a broad electron acceptor specificity and efficiently reduces substrates such as nitrofurazone, likely producing a two-electron transfer product | P17117 |
| **DegP** | DegP acts as a chaperone at low temperatures and converts to a serine protease (heat shock protein) under elevated temperatures to degrade misfolded proteins. | P0C0V0 |
| **ClpP** | ATP-dependent serine protease with chymotrypsin-like activity that mediates targeted degradation of misfolded or damaged proteins, playing a critical role in stress response. | P0A6G7 |
| **KatG** | Bifunctional enzyme with both catalase and broad-spectrum peroxidase activity. additionally exhibits NADH oxidase, isonicotinic acid hydrazide (INH) lyase, and isonicotinoyl-NAD synthase functions. | P13029 |
| **EntB** | Involved in the biosynthesis of the siderophore enterobactin (enterochelin), a trimeric macrocyclic lactone composed of N-(2,3-dihydroxybenzoyl)-L-serine residues. | P0ADI4 |
| **DacB** | Exhibits DD-carboxypeptidase and DD-endopeptidase activity but lacks transpeptidase function. Involved in cell wall remodeling and peptidoglycan metabolism. | P24228 |
| **CysE** | Catalyzes the initial step of cysteine biosynthesis by transferring an acetyl group from acetyl-CoA to L-serine, forming O-acetyl-L-serine. | P0A9D4 |
| **EntF** | Enzyme involved in the nonribosomal peptide synthesis of enterobactin, which is a macrocyclic trimeric lactone of N-(2,3-dihydroxybenzoyl)-serine | P11454 |
| **EntC** | Catalyzes the reversible isomerization of chorismate to isochorismate as part of the biosynthetic pathway for enterobactin, a siderophore composed of a trimeric lactone of N-(2,3-dihydroxybenzoyl)-L-serine. | P0AEJ2 |

[a] The protein function was obtained from UniProt [4].

[b] The UniProt accession code  for each protein [4].

**Table S7**. Biological processes associated with proteins involved in antibiotic resistance that increase in abundance during the exponential phase.

| | LOGARITMIC PHASE PROTEIN | | | | | | | | | | | | | |
|---|---|---|---|---|---|---|---|---|---|---|---|---|---|---|
| **Biological Process** | **CBAP** | **FeoB** | **EptA** | **IscR** | **PstB** | **EmrA** | **E_SOD** | **CusC** | **Lpp** | **DapE** | **PNPase** | **BamE** | **FabA** | **ArnA** |
| Guanosine nucleotides degradation III (*) | X | | | | | | | | | | | | | |
| Adenosine nucleotides degradation II (*) | X | | | | | | | | | | | | | |
| DNA damage response (GO:0006974) | | X | | | | | | | | | | | | |
| Iron ion import across plasma membrane (GO:0098711) | | X | | | | | | | | | | | | |
| Iron ion transport (GO:0006826) | | X | | | | | | | | | | | | |
| Lipopolysaccharide biosynthetic process (GO:0009103) | | | X | | | | | | | | | | | X |
| Lipid A biosynthetic process (GO:0009245) | | | X | | | | | | | | | | | X |
| Regulation of DNA-templated transcription (GO:0006355) | | | | X | | | | | | | | | | |
| Phosphate ion transport (GO:0006817) | | | | | X | | | | | | | | | |
| Phosphate ion transmembrane transport (GO:0035435) | | | | | X | | | | | | | | | |
| Monoatomic anion transmembrane transport (GO:0098656) | | | | | X | | | | | | | | | |
| Response to toxic substance (GO:0009636) | | | | | | X | | X | | | | | | |
| Bile acid and bile salt transport (GO:0015721) | | | | | | X | | | | | | | | |
| Response to steroid hormone (GO:0048545) | | | | | | X | | | | | | | | |
| Xenobiotic detoxification by transmembrane export across the plasma membrane (GO:1990961) | | | | | | X | | | | | | | | |
| Xenobiotic detoxification by transmembrane export across the cell outer membrane (GO:0140330) | | | | | | X | | | | | | | | |
| Superoxide metabolic process (GO:0006801) | | | | | | | X | | | | | | | |
| Removal of superoxide radicals (GO:0019430) | | | | | | | X | | | | | | | |
| Intracellular copper ion homeostasis (GO:0006878) | | | | | | | | X | | | | | | |
| Response to silver ion (GO:0010272) | | | | | | | | X | | | | | | |
| Detoxification of copper ion (GO:0010273) | | | | | | | | X | | | | | | |
| Response to copper ion (GO:0046688) | | | | | | | | X | | | | | | |
| Copper ion export (GO:0060003) | | | | | | | | X | | | | | | |
| Protein homotrimerization (GO:0070207) | | | | | | | | X | | | | | | |
| Monoatomic ion transport (GO:0006811) | | | | | | | | X | | | | | | |
| Copper ion transmembrane transport (GO:0035434) | | | | | | | | X | | | | | | |
| Transmembrane transport (GO:0055085) | | | | | | | | X | | | | | | |
| Silver ion transmembrane transport (GO:1902601) | | | | | | | | X | | | | | | |
| Lipid modification (GO:0030258) | | | | | | | | | X | | | | | |
| Periplasmic space organization (GO:0043580) | | | | | | | | | X | | | | | |
| FtsZ-dependent cytokinesis (GO:0043093) | | | | | | | | | | X | | | | |
| Lysine biosynthetic process via diaminopimelate (GO:0009089) | | | | | | | | | | X | | | | |
| Diaminopimelate biosynthetic process (GO:0019877) | | | | | | | | | | X | | | | |
| Response to heat (GO:0009408) | | | | | | | | | | | X | | | |
| RNA processing (GO:0006396) | | | | | | | | | | | X | | | |
| RNA catabolic process (GO:0006401) | | | | | | | | | | | X | | | |
| mRNA catabolic process (GO:0006402) | | | | | | | | | | | X | | | |
| Gram-negative-bacterium-type cell outer membrane assembly (GO:0043165) | | | | | | | | | | | | X | | |
| Protein insertion into membrane (GO:0051205) | | | | | | | | | | | | X | | |
| Fatty acid biosynthetic process (GO:0006633) | | | | | | | | | | | | | X | |
| Unsaturated fatty acid biosynthetic process (GO:0006636) | | | | | | | | | | | | | X | |
| UDP-D-xylose biosynthetic process (GO:0033320) | | | | | | | | | | | | | | X |
| UDP-4-deoxy-4-formamido-beta-L-arabinopyranose biosynthetic process (GO:2001315) | | | | | | | | | | | | | | X |
| Biosynthetic process (GO:0009058) | | | | | | | | | | | | | | X |
| Lipid A metabolic process (GO:0009245) | | | | | | | | | | | | | | X |
| Antibiotic Resistance | Aminoglycoside [5, 6] | Metronidazole [7] | Colistin [8-10] | Virulence Factor [11] | Virulence Factor [12] | Ampicillin [13] | Multidrug [14] | Carbapenem [15] | Antimicrobial peptides [16, 17] | Virulence Factor [18, 19] | Aminoglycoside [20] | Erythromycin [21] | Virulence Factor [22, 23] | Polymyxyn [24-26] |
| | Amphenicol [27] | | | | Penicillin [28] | Thiolactomycin [29] | | Tigecycline [15] | Vancomycin [30, 31] | | Ciprofloxacin [32] | | | |
| | Kanamycin [33] | | | | Vancomycin [34] | Multidrug [35] | | | | | Multidrug [36, 37] | | | |

The Gene Ontology (GO) code corresponding to the biological process in which each protein participates was obtained from the EcoCyc database [2, 3, 38]. The GO biological process for CBAP was not available in the EcoCyc database; instead, the information was retrieved from the UniProt database [4].

**Table S8**. Biological processes associated with proteins involved in antibiotic resistance that increase in abundance during the stationary phase.

| | STATIONARY PHASE PROTEIN | | | | | | | | | | | | |
|---|---|---|---|---|---|---|---|---|---|---|---|---|---|
| **Biological Process** | **HflC** | **NfsB** | **Mfd** | **AcrA** | **NfsA** | **DegP** | **ClpP** | **KatG** | **EntB** | **DacB** | **CysE** | **EntF** | **EntC** |
| Response to heat (GO:0009408) | X | | | | | X | X | | | | | | |
| 2,4,6-trinitrotoluene catabolic process (GO:0046256) | | X | | | | | | | | | | | |
| Transcription-coupled nucleotide-excision repair, DNA damage recognition (GO:0000716) | | | X | | | | | | | | | | |
| DNA repair (GO:0006281) | | | X | | | | | | | | | | |
| Transcription-coupled nucleotide-excision repair (GO:0006283) | | | X | | | | | | | | | | |
| Regulation of DNA-templated transcription (GO:0006355) | | | X | | | | | | | | | | |
| DNA damage response (GO:0006974) | | | X | | | | | | | | | | |
| Nucleotide-excision repair, preincision complex assembly (GO:0006294) | | | X | | | | | | | | | | |
| Response to xenobiotic stimulus (GO:0009410) | | | | X | | | | | | | | | |
| Response to toxic substance (GO:0009636) | | | | X | | | | | | | | | |
| Bile acid and bile salt transport (GO:0015721) | | | | X | | | | | | | | | |
| Alkane transport (GO:0015895) | | | | X | | | | | | | | | |
| Fatty acid transport (GO:0015908) | | | | X | | | | | | | | | |
| Xenobiotic transport (GO:0042908) | | | | X | | | | | | | | | |
| Xenobiotic detoxification by transmembrane export across the cell outer membrane (GO:0140330) | | | | X | | | | | | | | | |
| Transmembrane transport (GO:0055085) | | | | X | | | | | | | | | |
| Reduction of nitroaromatic compounds and quinones (*) | | | | | X | | | | | | | | |
| Oxidative stress response (*) | | | | | X | X | | X | | | | | |
| Interaction with tricarboxylic acid (TCA) cycle metabolites (*) | | | | | X | | | | | | | | |
| Protein folding (GO:0006457) | | | | | | X | | | | | | | |
| Proteolysis (GO:0006508) | | | | | | X | X | | | X | | | |
| Protein quality control for misfolded or incompletely synthesized proteins (GO:0006515) | | | | | | X | X | | | | | | |
| Response to temperature stimulus (GO:0009266) | | | | | | X | X | | | | | | |
| Chaperone-mediated protein folding (GO:0061077) | | | | | | X | | | | | | | |
| Response to radiation (GO:0009314) | | | | | | | X | | | | | | |
| Proteasomal protein catabolic process (GO:0010498) | | | | | | | X | | | | | | |
| Positive regulation of programmed cell death (GO:0043068) | | | | | | | X | | | | | | |
| Hydrogen peroxide catabolic process (GO:0042744) | | | | | | | | X | | | | | |
| Cellular response to hydrogen peroxide (GO:0070301) | | | | | | | | X | | | | | |
| Cellular oxidant detoxification (GO:0098869) | | | | | | | | X | | | | | |
| Enterobactin biosynthetic process (GO:0009239) | | | | | | | | | X | | | X | X |
| Peptidoglycan metabolic process (GO:0000270) | | | | | | | | | | X | | | |
| FtsZ-dependent cytokinesis (GO:0043093) | | | | | | | | | | X | | | |
| Regulation of cell shape (GO:0008360) | | | | | | | | | | X | | | |
| Peptidoglycan biosynthetic process (GO:0009252) | | | | | | | | | | X | | | |
| Cell division (GO:0051301) | | | | | | | | | | X | | | |
| Cell wall organization (GO:0071555) | | | | | | | | | | X | | | |
| Cysteine biosynthetic process from serine (GO:0006535) | | | | | | | | | | | X | | |
| Response to X-ray (GO:0010165) | | | | | | | | | | | X | | |
| Cysteine biosynthetic process (GO:0019344) | | | | | | | | | | | X | | |
| Amino acid activation for nonribosomal peptide biosynthetic process (GO:0043041) | | | | | | | | | | | | X | |
| Biosynthetic process (GO:0009058) | | | | | | | | | | | | X | X |
| Secondary metabolite biosynthetic process (GO:0044550) | | | | | | | | | | | | X | |
| Antibiotic Resistance | Aminoglycoside[39, 40] | Chloramphenicol[41, 42] | Fluoroquinolone[43] | Chloramphenicol[44] | 5-Nitrofuran derivates [45] | Virulence factor[46] | Daptomycin[47] | Isoniazid[48, 49] | Multidrug [50] | Penicillin[51, 52] | Mecillinam[53] | Virulence factor[54] | Virulence factor[55] |
| | | Nitrofurantoin[56, 57] | Multidrug[58 59] | Tigecycline [47] | Nitrofurantoin[56, 57 60 61] | | Multidrug[62, 63] | Virulence factor[64] | Virulence factor[65] | | Methicillin[66] | | |
| | | | | Multidrug[67-69] | | | | | | | Novobiocin[70] | | |
| | | | | | | | | | | | Multidrug[71] | | |

The Gene Ontology (GO) code corresponding to the biological process in which each protein participates was obtained from the EcoCyc database [2, 3, 38]. The GO biological process for NfsA was not available in the EcoCyc database; instead, the information was retrieved from the UniProt database [4].

[3] Gene Ontology, C., and Aleksander, S. A., and Balhoff, J., and Carbon, S., and Cherry, J. M., and Drabkin, H. J., and Ebert, D., and Feuermann, M., and Gaudet, P., and Harris, N. L., and Hill, D. P., and Lee, R., and Mi, H., and Moxon, S., and Mungall, C. J., and Muruganugan, A., and Mushayahama, T., and Sternberg, P. W., and Thomas, P. D., and Van Auken, K., and Ramsey, J., and Siegele, D. A., and Chisholm, R. L., and Fey, P., and Aspromonte, M. C., and Nugnes, M. V., and Quaglia, F., and Tosatto, S., and Giglio, M., and Nadendla, S., and Antonazzo, G., and Attrill, H., and Dos Santos, G., and Marygold, S., and Strelets, V., and Tabone, C. J., and Thurmond, J., and Zhou, P., and Ahmed, S. H., and Asanitthong, P., and Luna Buitrago, D., and Erdol, M. N., and Gage, M. C., and Ali Kadhum, M., and Li, K. Y. C., and Long, M., and Michalak, A., and Pesala, A., and Pritazahra, A., and Saverimuttu, S. C. C., and Su, R., and Thurlow, K. E., and Lovering, R. C., and

Logie, C., and Oliferenko, S., and Blake, J., and Christie, K., and Corbani, L., and Dolan, M. E., and Drabkin, H. J., and Hill, D. P., and Ni, L., and Sitnikov, D., and Smith, C., and Cuzick, A., and Seager, J., and Cooper, L., and Elser, J., and Jaiswal, P., and Gupta, P., and Jaiswal, P., and Naithani, S., and Lera-Ramirez, M., and Rutherford, K., and Wood, V., and De Pons, J. L., and Dwinell, M. R., and Hayman, G. T., and Kaldunski, M. L., and Kwitek, A. E., and Laulederkind, S. J. F., and Tutaj, M. A., and Vedi, M., and Wang, S. J., and D'Eustachio, P., and Aimo, L., and Axelsen, K., and Bridge, A., and Hyka-Nouspikel, N., and Morgat, A., and Aleksander, S. A., and Cherry, J. M., and Engel, S. R., and Karra, K., and Miyasato, S. R., and Nash, R. S., and Skrzypek, M. S., and Weng, S., and Wong, E. D., and Bakker, E., and Berardini, T. Z., and Reiser, L., and Auchincloss, A., and Axelsen, K., and Argoud-Puy, G., and Blatter, M. C., and Boutet, E., and Breuza, L., and Bridge, A., and Casals-Casas, C., and Coudert, E., and Estreicher, A., and Livia Famiglietti, M., and Feuermann, M., and Gos, A., and Gruaz-Gumowski, N., and Hulo, C., and Hyka-Nouspikel, N., and Jungo, F., and Le Mercier, P., and Lieberherr, D., and Masson, P., and Morgat, A., and Pedruzzi, I., and Pourcel, L., and Poux, S., and Rivoire, C., and Sundaram, S., and Bateman, A., and Bowler-Barnett, E., and Bye, A. J. H., and Denny, P., and Ignatchenko, A., and Ishtiaq, R., and Lock, A., and Lussi, Y., and Magrane, M., and Martin, M. J., and Orchard, S., and Raposo, P., and Speretta, E., and Tyagi, N., and Warner, K., and Zaru, R., and Diehl, A. D., and Lee, R., and Chan, J., and Diamantakis, S., and Raciti, D., and Zarowiecki, M., and Fisher, M., and James-Zorn, C., and Ponferrada, V., and Zorn, A., and Ramachandran, S., and Ruzicka, L., and Westerfield, M. (2023) The Gene Ontology knowledgebase in 2023, *Genetics 224*.

[4] UniProt, C. (2025) UniProt: the Universal Protein Knowledgebase in 2025, *Nucleic Acids Res 53*, D609-D617.

[5] Qin, S., Wang, Y., Zhang, Q., Chen, X., Shen, Z., Deng, F., Wu, C., and Shen, J. (2012) Identification of a novel genomic island conferring resistance to multiple aminoglycoside antibiotics in Campylobacter coli, *Antimicrob Agents Chemother 56*, 5332-5339.

[6] Efimochkina, N. R., Stetsenko, V. V., Bykova, I. V., Markova, Y. M., Polyanina, A. S., Aleshkina, A. I., and Sheveleva, S. A. (2018) Studying the Phenotypic and Genotypic Expression of Antibiotic Resistance in Campylobacter jejuni under Stress Conditions, *Bull Exp Biol Med 164*, 466-472.

[7] Veeranagouda, Y., Husain, F., Boente, R., Moore, J., Smith, C. J., Rocha, E. R., Patrick, S., and Wexler, H. M. (2014) Deficiency of the ferrous iron transporter FeoAB is linked with metronidazole resistance in Bacteroides fragilis, *J Antimicrob Chemother 69*, 2634-2643.

[8] Trebosc, V., Gartenmann, S., Totzl, M., Lucchini, V., Schellhorn, B., Pieren, M., Lociuro, S., Gitzinger, M., Tigges, M., Bumann, D., and Kemmer, C. (2019) Dissecting Colistin Resistance Mechanisms in Extensively Drug-Resistant Acinetobacter baumannii Clinical Isolates, *mBio 10*.

[9] Potron, A., Vuillemenot, J. B., Puja, H., Triponney, P., Bour, M., Valot, B., Amara, M., Cavalie, L., Bernard, C., Parmeland, L., Reibel, F., Larrouy-Maumus, G., Dortet, L., Bonnin, R. A., and Plesiat, P. (2019) ISAba1-dependent overexpression of eptA in clinical strains of Acinetobacter baumannii resistant to colistin, *J Antimicrob Chemother 74*, 2544-2550.

[10] Cervoni, M., Sposato, D., Lo Sciuto, A., and Imperi, F. (2023) Regulatory Landscape of the Pseudomonas aeruginosa Phosphoethanolamine Transferase Gene eptA in the Context of Colistin Resistance, *Antibiotics (Basel) 12*.

[11] Mettert, E. L., and Kiley, P. J. (2024) Fe-S cluster homeostasis and beyond: The multifaceted roles of IscR, *Biochim Biophys Acta Mol Cell Res 1871*, 119749.

[12] Yi, X., Xu, X., Qi, X., Chen, Y., Zhu, Z., Xu, G., Li, H., Kraco, E. K., Shen, H., Lin, M., Zheng, J., Qin, Y., and Jiang, X. (2022) Mechanisms Underlying the Virulence Regulation of Vibrio alginolyticus ND-01 pstS and pstB with a Transcriptomic Analysis, *Microorganisms 10*.

[13] Zhang, H., Ma, Y., Liu, P., and Li, X. (2016) Multidrug resistance operon emrAB contributes for chromate and ampicillin co-resistance in a Staphylococcus strain isolated from refinery polluted river bank, *Springerplus 5*, 1648.

[14] Heindorf, M., Kadari, M., Heider, C., Skiebe, E., and Wilharm, G. (2014) Impact of Acinetobacter baumannii superoxide dismutase on motility, virulence, oxidative stress resistance and susceptibility to antibiotics, *PLoS One 9*, e101033.

[15] Chen, D., Zhao, Y., Qiu, Y., Xiao, L., He, H., Zheng, D., Li, X., Yu, X., Xu, N., Hu, X., Chen, F., Li, H., and Chen, Y. (2019) CusS-CusR Two-Component System Mediates Tigecycline Resistance in Carbapenem-Resistant Klebsiella pneumoniae, *Front Microbiol 10*, 3159.

[16] Chang, T. W., Lin, Y. M., Wang, C. F., and Liao, Y. D. (2012) Outer membrane lipoprotein Lpp is Gram-negative bacterial cell surface receptor for cationic antimicrobial peptides, *J Biol Chem 287*, 418-428.

[17] Ebbensgaard, A., Mordhorst, H., Aarestrup, F. M., and Hansen, E. B. (2018) The Role of Outer Membrane Proteins and Lipopolysaccharides for the Sensitivity of Escherichia coli to Antimicrobial Peptides, *Front Microbiol 9*, 2153.

[18] Nocek, B., Starus, A., Makowska-Grzyska, M., Gutierrez, B., Sanchez, S., Jedrzejczak, R., Mack, J. C., Olsen, K. W., Joachimiak, A., and Holz, R. C. (2014) The dimerization domain in DapE enzymes is required for catalysis, *PLoS One 9*, e93593.

[19] Terrazas-Lopez, M., Gonzalez-Segura, L., Diaz-Vilchis, A., Aguirre-Mendez, K. A., Lobo-Galo, N., Martinez-Martinez, A., and Diaz-Sanchez, A. G. (2024) The three-dimensional structure of DapE from Enterococcus faecium reveals new insights into DapE/ArgE subfamily ligand specificity, *Int J Biol Macromol 270*, 132281.

[20] Fan, Z., Pan, X., Wang, D., Chen, R., Fu, T., Yang, B., Jin, Y., Bai, F., Cheng, Z., and Wu, W. (2021) Pseudomonas aeruginosa Polynucleotide Phosphorylase Controls Tolerance to Aminoglycoside Antibiotics by Regulating the MexXY Multidrug Efflux Pump, *Antimicrob Agents Chemother 65*.

[21] Ross, J. I., Eady, E. A., Cove, J. H., and Baumberg, S. (1995) Identification of a chromosomally encoded ABC-transport system with which the staphylococcal erythromycin exporter MsrA may interact, *Gene 153*, 93-98.

[22] Moynie, L., Leckie, S. M., McMahon, S. A., Duthie, F. G., Koehnke, A., Taylor, J. W., Alphey, M. S., Brenk, R., Smith, A. D., and Naismith, J. H. (2013) Structural insights into the mechanism and inhibition of the beta-hydroxydecanoyl-acyl carrier protein dehydratase from Pseudomonas aeruginosa, *J Mol Biol 425*, 365-377.

[23] Moynie, L., Hope, A. G., Finzel, K., Schmidberger, J., Leckie, S. M., Schneider, G., Burkart, M. D., Smith, A. D., Gray, D. W., and Naismith, J. H. (2016) A Substrate

Mimic Allows High-Throughput Assay of the FabA Protein and Consequently the Identification of a Novel Inhibitor of Pseudomonas aeruginosa FabA, *J Mol Biol 428*, 108-120.

[24] Gatzeva-Topalova, P. Z., May, A. P., and Sousa, M. C. (2004) Crystal structure of Escherichia coli ArnA (PmrI) decarboxylase domain. A key enzyme for lipid A modification with 4-amino-4-deoxy-L-arabinose and polymyxin resistance, *Biochemistry 43*, 13370-13379.

[25] Parulekar, R. S., Barage, S. H., Jalkute, C. B., Dhanavade, M. J., Fandilolu, P. M., and Sonawane, K. D. (2013) Homology modeling, molecular docking and DNA binding studies of nucleotide excision repair UvrC protein from M. tuberculosis, *Protein J 32*, 467-476.

[26] Mitchell, M. E., Gatzeva-Topalova, P. Z., Bargmann, A. D., Sammakia, T., and Sousa, M. C. (2023) Targeting the Conformational Change in ArnA Dehydrogenase for Selective Inhibition of Polymyxin Resistance, *Biochemistry 62*, 2216-2227.

[27] Qian, Y., Lai, L., Cheng, M., Fang, H., Fan, D., Zylstra, G. J., and Huang, X. (2024) Identification, characterization, and distribution of novel amidase gene aphA in sphingomonads conferring resistance to amphenicol antibiotics, *Appl Environ Microbiol 90*, e0151224.

[28] Engel, H., Mika, M., Denapaite, D., Hakenbeck, R., Muhlemann, K., Heller, M., Hathaway, L. J., and Hilty, M. (2014) A low-affinity penicillin-binding protein 2x variant is required for heteroresistance in Streptococcus pneumoniae, *Antimicrob Agents Chemother 58*, 3934-3941.

[29] Furukawa, H., Tsay, J. T., Jackowski, S., Takamura, Y., and Rock, C. O. (1993) Thiolactomycin resistance in Escherichia coli is associated with the multidrug resistance efflux pump encoded by emrAB, *J Bacteriol 175*, 3723-3729.

[30] Mathelie-Guinlet, M., Asmar, A. T., Collet, J. F., and Dufrene, Y. F. (2020) Lipoprotein Lpp regulates the mechanical properties of the E. coli cell envelope, *Nat Commun 11*, 1789.

[31] Wykes, H., Le, V. V. H., Olivera, C., and Rakonjac, J. (2023) When less is more: shortening the Lpp protein leads to increased vancomycin resistance in Escherichia coli, *J Antibiot (Tokyo) 76*, 746-750.

[32] Fan, Z., Chen, H., Li, M., Pan, X., Fu, W., Ren, H., Chen, R., Bai, F., Jin, Y., Cheng, Z., Jin, S., and Wu, W. (2019) Pseudomonas aeruginosa Polynucleotide Phosphorylase Contributes to Ciprofloxacin Resistance by Regulating PrtR, *Front Microbiol 10*, 1762.

[33] Gibreel, A., Skold, O., and Taylor, D. E. (2004) Characterization of plasmid-mediated aphA-3 kanamycin resistance in Campylobacter jejuni, *Microb Drug Resist 10*, 98-105.

[34] Bhardwaj, P., Islam, M. Z., Kim, C., Nguyen, U. T., and Palmer, K. L. (2021) ddcP, pstB, and excess D-lactate impact synergism between vancomycin and chlorhexidine against Enterococcus faecium 1,231,410, *PLoS One 16*, e0249631.

[35] Lomovskaya, O., and Lewis, K. (1992) Emr, an Escherichia coli locus for multidrug resistance, *Proc Natl Acad Sci U S A 89*, 8938-8942.

[36] McMurry, L. M., and Levy, S. B. (1987) Tn5 insertion in the polynucleotide phosphorylase (pnp) gene in Escherichia coli increases susceptibility to antibiotics, *J Bacteriol 169*, 1321-1324.

[37] Wu, N., Zhang, Y., Zhang, S., Yuan, Y., Liu, S., Xu, T., Cui, P., Zhang, W., and Zhang, Y. (2023) Polynucleotide Phosphorylase Mediates a New Mechanism of Persister Formation in Escherichia coli, *Microbiol Spectr 11*, e0154622.

[38] Moore, L. R., Caspi, R., Boyd, D., Berkmen, M., Mackie, A., Paley, S., and Karp, P. D. (2024) Revisiting the y-ome of Escherichia coli, *Nucleic Acids Res 52*, 12201-12207.

[39] Hinz, A., Lee, S., Jacoby, K., and Manoil, C. (2011) Membrane proteases and aminoglycoside antibiotic resistance, *J Bacteriol 193*, 4790-4797.

[40] Wessel, A. K., Yoshii, Y., Reder, A., Boudjemaa, R., Szczesna, M., Betton, J. M., Bernal-Bayard, J., Beloin, C., Lopez, D., Volker, U., and Ghigo, J. M. (2023) Escherichia coli SPFH Membrane Microdomain Proteins HflKC Contribute to Aminoglycoside and Oxidative Stress Tolerance, *Microbiol Spectr 11*, e0176723.

[41] Mullowney, M. W., Maltseva, N. I., Endres, M., Kim, Y., Joachimiak, A., and Crofts, T. S. (2022) Functional and Structural Characterization of Diverse NfsB Chloramphenicol Reductase Enzymes from Human Pathogens, *Microbiol Spectr 10*, e0013922.

[42] Crofts, T. S., Sontha, P., King, A. O., Wang, B., Biddy, B. A., Zanolli, N., Gaumnitz, J., and Dantas, G. (2019) Discovery and Characterization of a Nitroreductase Capable of Conferring Bacterial Resistance to Chloramphenicol, *Cell Chem Biol 26*, 559-570 e556.

[43] Han, J., Sahin, O., Barton, Y. W., and Zhang, Q. (2008) Key role of Mfd in the development of fluoroquinolone resistance in Campylobacter jejuni, *PLoS Pathog 4*, e1000083.

[44] Potrykus, J., Baranska, S., and Wegrzyn, G. (2002) Inactivation of the acrA gene is partially responsible for chloramphenicol sensitivity of Escherichia coli CM2555 strain expressing the chloramphenicol acetyltransferase gene, *Microb Drug Resist 8*, 179-185.

[45] Whiteway, J., Koziarz, P., Veall, J., Sandhu, N., Kumar, P., Hoecher, B., and Lambert, I. B. (1998) Oxygen-insensitive nitroreductases: analysis of the roles of nfsA and nfsB in development of resistance to 5-nitrofuran derivatives in Escherichia coli, *J Bacteriol 180*, 5529-5539.

[46] Cho, H., Choi, Y., Min, K., Son, J. B., Park, H., Lee, H. H., and Kim, S. (2020) Over-activation of a nonessential bacterial protease DegP as an antibiotic strategy, *Commun Biol 3*, 547.

[47] Xu, L., Henriksen, C., Mebus, V., Guerillot, R., Petersen, A., Jacques, N., Jiang, J. H., Derks, R. J. E., Sanchez-Lopez, E., Giera, M., Leeten, K., Stinear, T. P., Oury, C., Howden, B. P., Peleg, A. Y., and Frees, D. (2023) A Clinically Selected Staphylococcus aureus clpP Mutant Survives Daptomycin Treatment by Reducing Binding of the Antibiotic and Adapting a Rod-Shaped Morphology, *Antimicrob Agents Chemother 67*, e0032823.

[48] Brossier, F., Boudinet, M., Jarlier, V., Petrella, S., and Sougakoff, W. (2016) Comparative study of enzymatic activities of new KatG mutants from low- and high-level isoniazid-resistant clinical isolates of Mycobacterium tuberculosis, *Tuberculosis (Edinb) 100*, 15-24.

[49] Munir, A., Wilson, M. T., Hardwick, S. W., Chirgadze, D. Y., Worrall, J. A. R., Blundell, T. L., and Chaplin, A. K. (2021) Using cryo-EM to understand

antimycobacterial resistance in the catalase-peroxidase (KatG) from Mycobacterium tuberculosis, *Structure 29*, 899-912 e894.

[56] Wan, Y., Mills, E., Leung, R. C. Y., Vieira, A., Zhi, X., Croucher, N. J., Woodford, N., Jauneikaite, E., Ellington, M. J., and Sriskandan, S. (2021) Alterations in chromosomal genes nfsA, nfsB, and ribE are associated with nitrofurantoin resistance in Escherichia coli from the United Kingdom, *Microb Genom 7*.

[57] Zhang, X., Zhang, Y., Wang, F., Wang, C., Chen, L., Liu, H., Lu, H., Wen, H., and Zhou, T. (2018) Unravelling mechanisms of nitrofurantoin resistance and epidemiological characteristics among Escherichia coli clinical isolates, *Int J Antimicrob Agents 52*, 226-232.

[58] Lee, G. H., Jeong, J. Y., Chung, J. W., Nam, W. H., Lee, S. M., Pak, J. H., Choi, K. D., Song, H. J., Jung, H. Y., and Kim, J. H. (2009) The Helicobacter pylori Mfd protein is important for antibiotic resistance and DNA repair, *Diagn Microbiol Infect Dis 65*, 454-456.

[59] Ragheb, M. N., Thomason, M. K., Hsu, C., Nugent, P., Gage, J., Samadpour, A. N., Kariisa, A., Merrikh, C. N., Miller, S. I., Sherman, D. R., and Merrikh, H. (2019) Inhibiting the Evolution of Antibiotic Resistance, *Mol Cell 73*, 157-165 e155.

[60] Day, M. A., Jarrom, D., Christofferson, A. J., Graziano, A. E., Anderson, J. L. R., Searle, P. F., Hyde, E. I., and White, S. A. (2021) The structures of E. coli NfsA bound to the antibiotic nitrofurantoin; to 1,4-benzoquinone and to FMN, *Biochem J 478*, 2601-2617.

[61] Dulyayangkul, P., Sealey, J. E., Lee, W. W. Y., Satapoomin, N., Reding, C., Heesom, K. J., Williams, P. B., and Avison, M. B. (2024) Improving nitrofurantoin resistance prediction in Escherichia coli from whole-genome sequence by integrating NfsA/B enzyme assays, *Antimicrob Agents Chemother 68*, e0024224.

[62] Frees, D., Gerth, U., and Ingmer, H. (2014) Clp chaperones and proteases are central in stress survival, virulence and antibiotic resistance of Staphylococcus aureus, *Int J Med Microbiol 304*, 142-149.

[63] Zheng, J., Wu, Y., Lin, Z., Wang, G., Jiang, S., Sun, X., Tu, H., Yu, Z., and Qu, D. (2020) ClpP participates in stress tolerance, biofilm formation, antimicrobial tolerance, and virulence of Enterococcus faecalis, *BMC Microbiol 20*, 30.

[64] Escalante, P., McKean-Cowdin, R., Ramaswamy, S. V., Williams-Bouyer, N., Teeter, L. D., Jones, B. E., and Graviss, E. A. (2013) Can mycobacterial katG genetic changes in isoniazid-resistant tuberculosis influence human disease features?, *Int J Tuberc Lung Dis 17*, 644-651.

[65] Amaretti, A., Righini, L., Candeliere, F., Musmeci, E., Bonvicini, F., Gentilomi, G. A., Rossi, M., and Raimondi, S. (2020) Antibiotic Resistance, Virulence Factors, Phenotyping, and Genotyping of Non-Escherichia coli Enterobacterales from the Gut Microbiota of Healthy Subjects, *Int J Mol Sci 21*.
